# How good are predictions of the effects of selective sweeps on levels of neutral diversity?

**DOI:** 10.1101/2020.05.27.119883

**Authors:** Brian Charlesworth

## Abstract

Selective sweeps are thought to play a significant role in shaping patterns of variability across genomes; accurate predictions of their effects are, therefore, important for understanding these patterns. A commonly used model of selective sweeps assumes that alleles sampled at the end of a sweep, and that fail to recombine with wild-type haplotypes during the sweep, coalesce instantaneously, leading to a simple expression for sweep effects on diversity. It is shown here that there can be a significant probability that a pair of alleles sampled at the end of a sweep coalesce during the sweep before a recombination event can occur, reducing their expected coalescent time below that given by the simple approximation. Expressions are derived for the expected reductions in pairwise neutral diversities caused by both single and recurrent sweeps in the presence of such within-sweep coalescence, although the effects of multiple recombination events during a sweep are only treated heuristically. The accuracies of the resulting expressions were checked against the results of simulations. For even moderate ratios of the recombination rate to the selection coefficient, the simple approximation can be substantially inaccurate. The selection model used here can be applied to favorable mutations with arbitrary dominance coefficients, to sex-linked loci with sex-specific selection coefficients, and to inbreeding populations. Using the results from this model, the expected differences between the levels of variability on X chromosomes and autosomes with selection at linked sites are discussed, and compared with data on a population of *Drosophila melanogaster*.

---

Maynard Smith and Haigh (1974) introduced the concept of hitchhiking into population genetics, showing how the spread of a favorable mutation reduces the level of neutral variability at a linked locus. Nearly twenty years later, it was shown that selection against recurrent deleterious mutations can also reduce variability, by the hitchhiking process known as background selection (Charlesworth *et al*. 1993). It is, therefore, preferable to use the term “selective sweep” (Berry and Kreitman 1993) for the hitchhiking effects of favorable mutations. There is now a large theoretical and empirical literature on both types of hitchhiking, recently reviewed by Walsh and Lynch (2018) and Stephan (2019). With sufficiently weak selection, recurrent partially recessive deleterious mutations can also increase, rather than reduce, variability at linked sites, because fluctuations in their frequencies due to genetic drift create associative overdominance (Zhao and Charlesworth 2016; Becher *et al*. 2020; Gilbert *et al*. 2020). The expected effects of all three processes on pairwise diversity at a neutral site can be described by the same general formula (Zhao and Charlesworth 2016), which is a consequence of the Price-Robertson equation for the change in the mean of a trait induced by its additive genetic covariance with fitness (Robertson 1968; Price 1970).

These theoretical studies have provided the basis for methods for inferring the nature and parameters of selection from population genomic data, recently reviewed by Booker *et al*. (2017). Several recent studies have concluded that the level of DNA sequence variability in a species is often much smaller than would be expected in the absence of selection (Corbett-Detig *et al*. 2015; Elyashiv *et al*. 2016; Campos *et al*. 2017; Comeron 2017), especially for synonymous sites in coding sequences, reflecting both the effects of both selective sweeps and background selection (BGS). However, estimates of the parameters involved differ substantially among different studies. There is also an ongoing debate about the extent to which the level of genetic variability in a species is controlled by classical genetic drift, reflecting its population size, or by the effects of selection in removing variability. The possibility that the effects of selective sweeps dominate over drift was originally raised by Maynard Smith and Haigh (1974), and later advocated by (Kaplan *et al*. (1989) and Gillespie (2002); see Kern and Hahn (2018) and Jensen *et al*. (2019) for recent discussions of this question.

The model of Maynard Smith and Haigh (1974) assumed that the trajectory of the selectively favored allele was purely deterministic. Kaplan *et al*. (1989) developed a representation of the dual processes of recombination and coalescence during a sweep, which allowed for stochastic effects on the frequency of the selected allele when it is either rare or very common. This approach enabled calculations of the effect of a sweep on both pairwise diversity and the site frequency spectrum, but did not provide simple formulae. Explicit formulae for the effect of a sweep on pairwise diversity that removed the assumption of a purely deterministic trajectory were derived by Stephan *et al*. (1992) using diffusion equations. Barton (1998, 2000) developed an alternative approach using a combination of branching processes and diffusion equations, from which the properties of a post-sweep sample of *n* alleles could be calculated. Kaplan *et al*. (1989), Stephan *et al*. (1992), Wiehe and Stephan (1993), Barton (2000), Kim and Stephan (2000) and Gillespie (2002) also analyzed the effects of recurrent selective sweeps, treating coalescent events caused by classical genetic drift and by sweeps as competing exponential processes. All of these approaches assumed either a haploid population or an autosomal locus with semi-dominant fitness effects.

A great simplification in such calculations was achieved by the following approximation, proposed by Barton (1998, 2000) and extended by Durrett and Schweinsberg (2004) – see also Coop and Ralph (2012). This approach is based on two assumptions. The first is that the fixation of a favorable mutation happens so fast that non-recombinant alleles at a linked neutral site, sampled after the completion of the sweep, coalesce at such a high rate that their coalescence time is negligible relative to that under neutrality. The second is that linkage is sufficiently tight that at most a single recombination event occurs during the sweep, placing a neutral site onto a wild-type background with which it remains associated throughout the sweep. These assumptions mean that the gene genealogy for a set of alleles sampled immediately after a sweep, and that failed to recombine onto the wild-type background, has a “star-like” shape. The reduction in diversity and site frequency spectrum at the neutral site can then be calculated in a straightforward fashion (Barton 2000; Durrett and Schweinsberg 2004; Kim 2006; Weissman and Barton 2012; Coop and Ralph 2012). This approximation provides the basis for detecting recent sweeps in the popular programs SweepFinder (Nielsen *et al*. 2005) and Sweed (Pavlidis *et al*. 2013). It can readily be incorporated into models of recurrent selective sweeps (Barton 2000; Weissman and Barton 2012; Berg and Coop 2015; Elyashiv *et al*. 2016; Campos *et al*. 2017; Campos and Charlesworth 2019), which has stimulated the development of methods for estimating the parameters of recurrent sweeps from population genomic data (Elyashiv *et al*. 2016; Campos *et al*. 2017; Campos and Charlesworth 2019).

This approach is likely to be accurate for favorable mutations that are sufficiently strongly selected that their time to fixation is short compared with the expected neutral coalescent time of 2*N_e_* generations (where *N_e_* is the effective population size), especially when the ratio of the recombination rate to the selection coefficient is small. Recent population genomic analyses suggest, however, that there may be important contributions from relatively weakly selected favorable mutations, which can take as long as 10% or more of the neutral coalescent time to become fixed (Sella *et al*. 2009; Keightley *et al*. 2016; Chen *et al*. 2020). In such cases, the time to coalescence during the sweep cannot necessarily be neglected, and the assumption that a pair of non-recombined alleles are identical in state leads to an overestimate of diversity at the end of the sweep, especially with very low rates of recombination. In contrast, coalescence during the sweep competes with recombination, so that calculating the probability that one of a pair of alleles recombines onto the wild-type background without including the probability that they have escaped prior coalescence underestimates the effect of a sweep (Barton 1998). More generally, when the assumption that the duration of a sweep is negligible compared with the neutral coalescent time is invalid, the mean coalescent time of a pair of alleles cannot accurately be calculated simply from the probability that they escape recombination onto the wild-type background.

The present paper describes a general analytical model of selective sweep effects on the mean time to coalescence of a pair of alleles at a linked neutral locus (which determines the expected pairwise neutral diversity), for the case of weak selection at a single locus, where the selection coefficient is sufficiently small that a differential equation can used instead of a difference equation. This is based on a recent study of the expected time to fixation of a favorable mutation in a single population (Charlesworth 2020), which provided a general framework for analyzing both autosomal and sex-linked inheritance with arbitrary levels of inbreeding and dominance.

The resulting formulae, which include a heuristic treatment of multiple recombination events, enable predictions of the effects on diversity of both a single sweep and recurrent selective sweeps, and allow for the action of BGS as well as sweeps. Hartfield and Bataillon (2020) have recently presented similar results for an autosomal locus with coalescence during a sweep, in the case of a single sweep in the absence of BGS, but without modelling multiple recombination events. Only hard sweeps will be considered here, although it is straightforward to extend the models to soft sweeps by the approach of Berg and Coop (2015) and Hartfield and Bataillon (2020). The validity of the approximations is tested against computer simulations, including those of Campos and Charlesworth (2019) and Hartfield and Bataillon (2020). For the sake of brevity, these papers will be referred to as CC and HB, respectively.

## Methods

### Simulating the effect of a single sweep

The algorithm described by Equations 27 of Tajima (1990) was used to calculate the effects of a sweep on pairwise diversity at a neutral locus with an arbitrary degree of linkage to a selected locus with two alleles, A_1_ and A_2_, where A_2_ is the selectively favored allele. A Wright-Fisher population with constant size *N* was assumed. The equations provide three coupled, forward-in-time recurrence relations for the expected diversities at the neutral locus for pairs of alleles carrying either A_1_ or A_2_, and for the divergence between A_1_ and A_2_ alleles. These are conditioned on a given generation-by-generation trajectory of allele frequencies at the selected locus, and assume an infinite sites model of mutation and drift (Kimura 1971).

The initial conditions for a simulation run were that a single A_2_ allele was introduced into the population, with zero expected pairwise diversity at the associated neutral locus; the expected pairwise diversity among A_1_ alleles and the divergence between A_1_ and A_2_ were equal to those for an equilibrium population in the absence of selection, *θ* = 4*Nu*, where *u* is the neutral mutation rate. Since only diversities relative to *θ* are of interest here, *θ* was set to 0.001 in order to satisfy the infinite sites assumption for the neutral locus. The expected change in the frequency *q* of A_2_ in a given generation for an assigned selection model was calculated using the standard discrete-generation selection formulation (see the section Theoretical Results for details of the models of selection). Binomial sampling using the frequency after selection and 2*N* as parameters was used to obtain the value of *q* in the next generation. Equations 27 of Tajima (1990) were applied to the old value of *q* in order to obtain the state of the neutral locus in the new generation.

This procedure was repeated generation by generation until A_2_ was lost or fixed; only runs in which A_2_ was lost were retained, and the value of the pairwise diversity among A_2_ alleles at the time of its fixation was determined. This gives the expected diversity after a sweep conditional on a given trajectory, so that an estimate of the overall expected diversity relative to *θ* can be found by taking the mean over a large number of replicate simulations. It was found that 100 replicates were sufficient to produce a standard error of 2% or less of the mean. The value of *N* was chosen so that the selection coefficient *s* for a given value of the scaled selection parameter *γ* = *2Ns* was sufficiently small that terms of order *s*^2^ could be neglected, to satisfy the assumptions of the model described in the Theoretical Results section.

### Recurrent sweeps: simulation methods

For checking the theoretical predictions concerning recurrent sweeps, the simulation results described in CC were used. These involved groups of linked autosomal genes separated by 2 kilobases of selectively neutral intergenic sequence, with all UTR sites and 70% of nonsynonymous (NS) sites subject to both positive and negative selection, and the same selection parameters for 5’and 3’ UTRs (see Figure 1 of CC). There were 5 exons of 300 basepairs (bp) each, interrupted by 4 introns of 100bp. The lengths of the 5’and 3’ UTRs were 190bp and 280bp, respectively. The selection coefficients for favorable and deleterious mutations at the NS and UTR sites, and the proportions of mutations at these sites that were favorable, were chosen to match the values inferred by Campos *et al*. (2017) from the relation between the synonymous diversity of a gene and its rate of protein sequence evolution. Both favorable and deleterious mutations were assumed to be semidominant.

**Figure 1.**
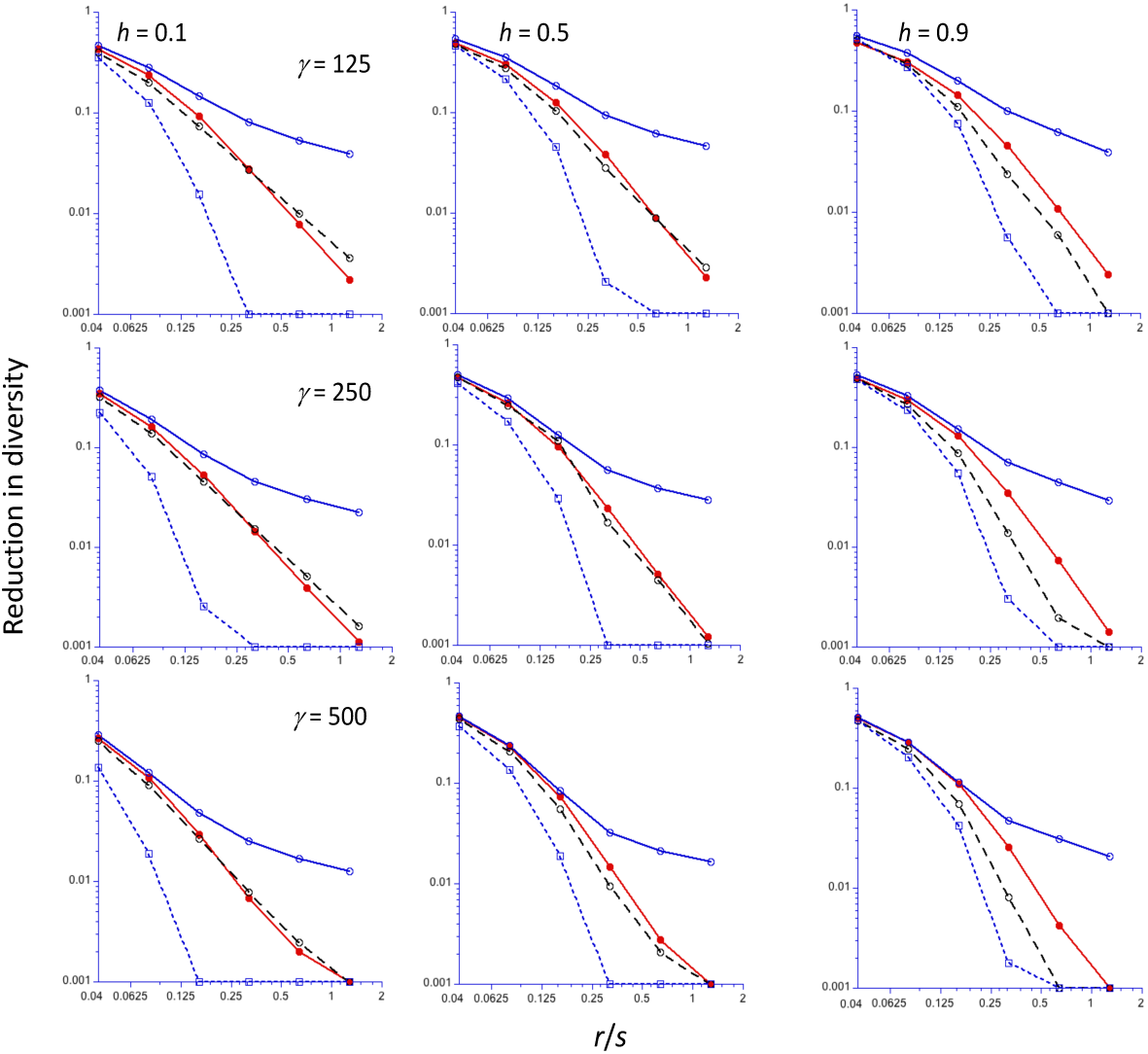
The reduction in diversity (relative to the neutral value) at the end of a sweep for an autosomal locus (Y-axis, log_10_ scale), as a function of the ratio of the frequency of recombination (*r*) to the selection coefficient for homozygotes (*s*) (X-axis, log_2_ scale). A population size of 5000 is assumed, with three different values of the scaled selection coefficient (*γ* = 2*N_e_s*): 125 (top panel), 250 (middle panel) and 500 (bottom panel), and three different values of the dominance coefficient (*h*), increasing from left to right. The filled red circles are the mean values from computer simulations, using the algorithm of Tajima (1990); the open filled blue circles and black circles are the *C1* and *C2* predictions, respectively; the open blue squares are the *NC* predictions. Values of the reduction in diversity less than 0.001 have been reset to 0.001.

Five different rates of reciprocal crossing over (CO) were used to model recombination, which were chosen to be multiples of the approximate standard autosomal recombination rate in *Drosophila melanogaster*, adjusted by a factor of ½ to take into account the absence of recombinational exchange in males (Campos *et al*. 2017): 0.5 x 10^-8^, 1 x 10^-8^, 1.5 x 10^-8^, 2 x 10^-8^ and 2.5 x 10^-8^ cM/Mb, respectively, where 10^-8^ is the mean rate across the genome.

The simulations were run with and without BGS acting on both NS and UTR sites, and with and without non-crossover associated gene conversion events. Cases with gene conversion assumed a rate of initiation of conversion events of 1 x 10^-8^cM/Mb for autosomes (after correcting for the lack of gene conversion in males), and a mean tract length of 440 bp, with an exponential distribution of tract lengths.

### Recurrent sweeps at multiple sites: numerical predictions based on analytical formulae

A single gene is considered in the analytical models, so that a linear genetic map can be assumed, because there is a negligible frequency of double crossovers. The CO contribution to the frequency of recombination between a pair of sites separated by *z* basepairs is *r_c_z*, where *z* is the physical distance between the neutral and selected sites and *r_c_* is the CO rate CO per bp.

An important point regarding the cases with gene conversion should be noted here. CC stated that, because the simulation program they used (SLiM 1.8) modeled gene conversion by considering only events that are initiated on one side of a given nucleotide site, the rate of initiation of a gene conversion tract covering this site is one-half of that used in the standard formula for the frequency of recombination caused by gene conversion; see Equation 1 of Frisse *et al*. (2001). However, this statement is incorrect, because it overlooks the fact that the standard model of gene conversion assumes that there are equal probabilities of a tract moving towards and away from the site. If tracts are constrained to move in one direction, the net probability that a tract started at a random point moves towards a given site is the same as in the standard formula, for a given probability of initiation of a tract.

Since no derivation of the formula of Frisse *et al*. (2001) appears to have been given, one is provided in Section S1 of File S1, which makes this point explicit (Equation S5 is equivalent to the formula in question). Gene conversion tract lengths are assumed to be exponentially distributed, with a mean tract length of *d_g_*, and a probability of initiation *r_g_*. It follows that the effective rates of initiation of gene conversion events (*r_g_*) used in the theoretical calculations in CC should have been twice the values that were used. Diversity values were thus under-estimated by these calculations, because there was more recombination than was included in the predictions. The correct theoretical results for sweep effects are presented here.

The effects of selective sweeps on neutral sites within a gene were obtained by summing the expected effects of substitutions at each NS and UTR site in the gene on a given neutral site (synonymous site), assuming that every third basepair in an exon is a neutral site, with the other two (NS) sites being subject to selection, as described by Campos *et al*. (2017). This differs from the SLiM procedure of randomly assigning selection status to exonic sites, with a probability *p_s_* of being under selection (*p_s_* = 0.7 in the simulations used in CC). To correct for this, the overall rate of NS substitutions per NS site was adjusted by multiplication by 0.7 x 1.5. Furthermore, to correct for the effects of interference among co-occurring favourable mutations in reducing their probabilities of fixation, their predicted rates of substitution were multiplied by a factor of 0.95, following the procedure in CC.

In order to speed up the computations, mean values of the variables used to calculate the effects of sweeps on neutral diversity were calculated by thinning the neutral sites by considering only a subset of them, starting with the first codon at the 5’end of the gene. For the results reported here, 10% of all neutral sites were used to calculate the values of the variables. Comparisons with results from using all sites showed a negligible effect of using this thinning procedure.

Background selection effects on diversity for autosomes and X chromosomes for genes in regions with different CO rates were calculated as described in sections S9 and S10 of File S1 of CC, which included estimates of the effects of BGS caused by selectively constrained non-coding sequences as well as coding sequences, derived from (Charlesworth 2012). If gene conversion was absent, the correction factors for gene conversion used to calculate these effects were omitted.

No new data or reagents were generated by this research. Details of some of the mathematical derivations are described in the Supplementary Information, File S1. The codes for the computer programs used to implement the analytical models described below will be made available in the Supplementary Information, File S2, on Figshare on acceptance. The detailed statistics for the results of the computer simulations were provided in Files S2-S3 of Campos and Charlesworth (2019).

## The effect of a single sweep on expected nucleotide site diversity

### Theoretical results

The aim of this section is obtain an expression for the mean coalescent time at a neutral site linked to a selected locus, at the time of fixation of the selectively favored allele; under the infinite sites model, this yields the expected pairwise diversity at the neutral site. All times are expressed on the coalescent timescale of 2*N_e_* generations, where *N_e_* is the neutral effective population size for the genetic system under consideration (autosomal or X-linked loci, random mating or partial inbreeding). If we use *N_e_*_0_ to denote the value of *N_e_* for a randomly mating population with autosomal inheritance, *N_e_* for a given genetic system can be written as *kN_e_*_0_, where *k* depends on the details of the system in question (Wright 1931,1969; Crow and Kimura 1970; Charlesworth and Charlesworth 2010). For example, with an autosomal locus in a partially inbreeding population with Wright’s fixation index *F* > 0, we have *k* ≈1/(1+*F*) under a wide range of conditions (Pollak 1987; Nordborg 1997; Laporte and Charlesworth 2002). In addition, following Kim and Stephan (2000) and CC, if BGS is operating, it is assumed that, for purely neutral processes, *N_e_* can replaced by the quantity *B*_1_*N_e_*, where *B*_1_ measures the effect of BGS on the mean neutral coalescent time of a pair of alleles. The effect of BGS on the fixation probabilities of favorable mutations is likely to be somewhat less than that for neutral processes, so that a second coefficient,*B*_2_, should ideally be used as a multiplier of *N_e_*, where *B*_2_ = *B*_1_/λ (λ ≤ 1). As discussed in CC, *B*_1_ can be determined analytically for a given genetic model, whereas *B*_2_ usually requires simulations, so it is often more convenient to use *B*_1_ for both purposes, although this procedure introduces some inaccuracies.

As has been discussed in previous treatments of sweeps, there are two stochastic phases during the spread of a favorable mutation, A_2_, in competition with a wild-type allele, A_1_. A detailed analysis of these stochastic phases for the general model of selection used here is given by Charlesworth (2020). In the first phase, the frequency of A_2_ is so low that it is subject to random fluctuations that can lead to the loss of A_2_ from the population. Provided that the product of *N_e_* and the selection coefficient for homozygotes for the favorable allele (*s*) is >> 1, a mutation that survives this phase will enter the deterministic phase, where it has a negligible probability of loss, and in which its trajectory of allele frequency change is well approximated by the deterministic selection equation (Equation 6 below). When A_2_ reaches a frequency close to 1, A_1_ is now vulnerable to stochastic loss, so that there is a second stochastic phase. Formulae for the frequencies of A_2_ at the boundaries of the two stochastic phases, *q*_1_ and *q*_2_,,are given by Charlesworth (2020), together with expressions for the durations of the stochastic and deterministic phases. For mutations with intermediate levels of dominance, *q*_1_, 1 – *q*_2_ and the durations of the two stochastic phases are all of the order of 1/(2*N_e_s*), measured on the coalescent timescale of 2*N_e_* generations.

If *q*_2_ is close to 1, A_2_ has only a small chance of encountering an A_1_ allele, so that there is a negligible chance that a neutral site in a haplotype carrying A_2_ will recombine onto an background recombination during the final stochastic phase. In addition, the rate of coalescence within haplotypes carrying A_2_ is then close to the neutral value, and so does not greatly affect the mean time to coalescence of a pair of alleles sampled after the end of the sweeep compared with neutral expectation. Under these conditions, the second stochastic phase has little effect on the mean coalescent time of the alleles compared with neutral expectation. Provided that the duration of the first stochastic phase on the coalescent time scale is << 1 (i.e. *q*_1_ is close to 0), this phase will also have a minimal impact on the mean coalescent time of such a pair of alleles. Accurate approximations for the effect of a single sweep on diversity can, therefore, usually be obtained by treating the the beginning and end of the deterministic phase as equivalent to that for the sweep as a whole, as discussed by Charlesworth (2020).

The general framework presented in HB can then be used to determine the effect of a sweep on pairwise diversity, extended to include a more general model of selection as well as the possibility of BGS effects, and using analytical expressions for probabilities of coalescence and recombination during the sweep rather than numerical evaluations. This approach assumes that all evolutionary forces are weak (i.e., second order terms in changes in allele frequencies and linkage disequilibrium can be neglected), so that a continuous time scale approximation to a discrete generation model can be applied.

Let *T_d_* be the duration of the deterministic phase, defined as the period between frequencies *q*_1_ and *q*_2_ as given by Charlesworth (2020). With BGS, the terms in *N_e_* in the relevant expressions are each to be multiplied by *B*_2_, as was done in CC. For a pair of haplotypes that carry the favorable allele A_2_ at the end of the sweep, the rate of coalescence at a time *T* back from this time point is [*B*_1_*q*(*T*)]^−1^, where *q*(*T*) is the frequency of A_2_ at time *T*. The rate at which a linked neutral site recombines from A_2_ onto the wild-type background at time *T* is *ρ*[1 – *q*(*T*)] =*ρp*(*T*), where *ρ* = 2*N_e_r* is the scaled recombination rate and *r* is the absolute recombination rate between the selected and neutral loci. With inbreeding and/or sex-linkage, *r* differs from its random mating autosomal value, *r*_0_, such that *r* = *cr*_0_, where *c* is a function of the genetic system and mating system. For example, with autosomal inheritance with partial inbreeding, *c* ≈ 1 – 2*F +ϕ*, where *ϕ* is the joint probability of identity by descent at a pair of neutral loci (Roze 2009; Hartfield and Bataillon 2020). Unless both *r*_0_ and 1 – *F* are sufficiently large that their second-order terms cannot be neglected, we have *c* ≈ 1 – *F* (Nordborg 1997; Charlesworth and Charlesworth 2010, p.381). The exact value of *ϕ* is determined by the mating system; in the case of self-fertilization, Equation 1 of HB gives an expression for *ϕ* as a function of *r*_0_ and the rate of self-fertilization, which is used in the calculations presented here.

Under these assumptions, the probability density function (p.d.f.) for a coalescent event at time *T* for a pair of alleles sampled at the end of the sweep is:

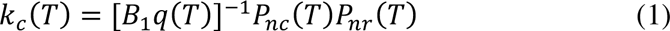

where *P_nc_*(*T*) is the probability of no coalescence by time*T* in the absence of recombination, and *P_nr_*(*T*) is the probability that neither allele has recombined onto the wild-type background by time*T*, in the absence of coalescence.

Similarly, the p.d.f. for the event that one of the two sampled haplotypes recombines onto the wild-type background at time *T* (assuming that *r* is sufficiently small that simultaneous recombination events can be ignored) is given by:

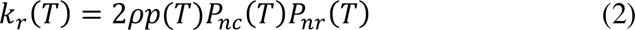

We therefore have:

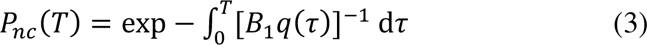

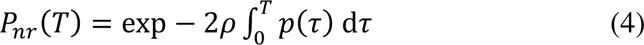

The net probability that the pair of sampled alleles coalesce during the deterministic phase of the sweep is given by:

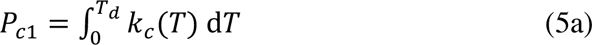

If it is assumed that haplotypes that have neither recombined nor coalesced during the sweep coalesce with probability one at the start of the sweep, there is an additional contribution to the coalescence probability, given by:

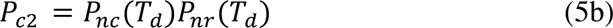

The net probability of coalescence caused by the sweep is thus:

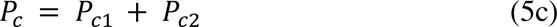

These equations are simple in form, but getting explicit formulae is made difficult by the non-linearity of the equation for the rate of change of *q* under selection. Following Charlesworth (2020), for the case of weak selection (when terms of order *s*^2^ can be ignored) we can write the forward-in-time selection equation as:

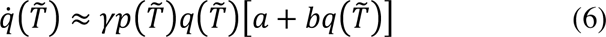

where tildes are used to denote time measured from the start of the sweep; γ = 2*N_e_s* is the scaled selection coefficient for A_2_A_2_, assigning a fitness of 1 tο A_1_A_1_ and an increase in relative fitness of *s* to A_2_A_2_. Here, *a* and *b* depend on the dominance coefficient *h* and fixation index *F*, the genetic and mating systems, and the sex-specificity of fitness effects (Glémin 2012; Charlesworth 2020). For example, for an autosomal locus, the weak selection approximation gives *a* = *F* + (1 – *F*)*h* and *b* = (1 – *F*)(1 – 2*h*).

For *a* > 0 and *a* + *b* > 0, corresponding to intermediate levels of dominance, integration of Equation 6 yields the following expression for the expectation of the duration of the deterministic phase, *T_d_* (Charlesworth 2020):

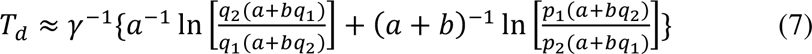

Similar expressions are available for the cases when *a* = 0 (complete recessivity) or *a* + *b* = 0 (complete dominance), as described by Charlesworth (2020).

Using Equation 6, we can write *T* as a monotonic function of *q*, *T*(*q*), substituting *q* for *T* and using the relation 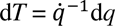. Equations 5a, 3 and 4 then become:

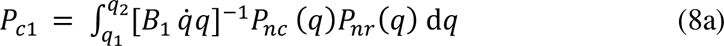

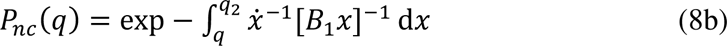

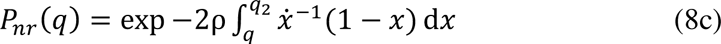

Substituting *q*_1_ for *q* in Equations 5b and 5c, Equation 5b can be written as:

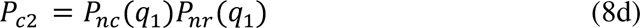

The net expected pairwise coalescence time associated with the sweep, *T_s_*, includes a contribution from the case when no coalescence occurs until the start of the sweep, given by the product of *P_c_*_2_ and *T_d_*, and a contribution from coalescent events that occur during the sweep, denoted by *T_c_*. We have:

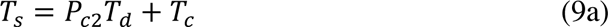

where

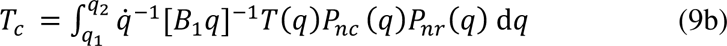

and *T*(*q*) is the time to coalescence at frequency *q* of A_2_, given by Equations A1.

#### Results with only a single recombination event

The possibility of recombination back onto the background of A_2_, examined in CC, is ignored for the present, as is the possibility of a second recombination event from A_2_ onto A_1_. From Equation 2, the probability of at least one recombination event is given by:

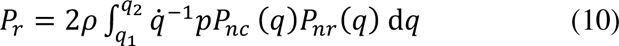

Using Equations 6, 10a and A1-A3, *P_r_* can be expressed explicitly in terms of *ρ*, *γ*, *a* and *b*, but has to be evaluated numerically.

The net expected pairwise coalescence time in the presence of BGS under this set of assumptions is given by *B*_1_*P_r_* +*T_s_*. Under the infinite sites model (Kimura 1971), the expected reduction in pairwise nucleotide site diversity for alleles sampled at the end of the sweep, relative to its value in the absence of selection (*θ*), is given by:

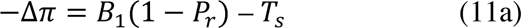

Equation 9 of HB for the case of a hard sweep is equivalent to Equation 11a without the term in *T_s_*. In addition, if *T_s_* and the probability of coalescence during the sweep are both negligible, it is easily seen that *P_r_* ≈ 1 – *P_nr_*(*T_d_*), yielding result for the star phylogeny approximation (Barton 1998, 2000; Durrett and Schweinsberg 2004; Weissman and Barton 2012):

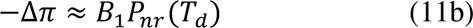

Explicit formulae for the components of the above equations, using Equation 6, are given in the Appendix. In the case of an autosomal locus with random mating and semi-dominant selection (*h* = 0.5), substitution of these formulae into Equation 11c yield the following convenient formula, which has been used in inferences from population genomic data, as mentioned in the introduction section:

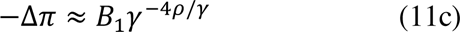

#### The importance of coalescence during a sweep

These results bring out the potential importance of considering coalescence during a sweep, as opposed to the coalescence of non-recombined alleles at the start of a sweep. Consider the case with incomplete dominance (*a* ≠ 0). The probability of no coalescence during the sweep conditional on no recombination, *P_nc_*(*q*_1_), is given by Equation A2a with *q* = *q*_1_, where *q*_1_ ≈ (2*aγ*)^−1^ (Charlesworth 2020). Somewhat surprisingly, for large γ this expression becomes independent of *a* and *γ*, provided that *a* ^−2^ >> *γ,* and approaches *e* ^−2^ ≈ 0.135, so that the probability of coalescence during a sweep in the absence of recombination is approximately 0.865 (see the Appendix). With low rates of recombination, there is thus a high probability of coalescence during the sweep itself, in contrast to what is assumed in Equations 11b and 11c. If such a coalescent event is not preceded by a recombination event, the mean coalescent time will thus be smaller than predicted by these Equations.

This raises the question of the magnitude of *T_s_* in the more exact treatment. While Equation 9 can only be evaluated exactly by numerical integration, a rough estimate of *T_s_* for the case of no recombination can be obtained as follows (this is the maximum value, as the terms involving the probability of no recombination must decrease with the frequency of recombination). By the above result for *P_nc_*(*q*_1_), the first term in Equation 9 is approximately *e* ^−2^*T_d_*. The second term is equivalent to the mean coalescent time associated with events during the sweep; by the argument presented in section S3 of File S1 in CC, this is approximately equal to the harmonic mean of 1/*q* between *q*_1_ and *q*_2_. Equation S10 of CC for this quantity can be generalized as shown in the Appendix, with the result that the expected coalescent time associated with the sweep (*T_c_*) is approximately ½*T_d_* for large *γ*, yielding *T_s_* ≈ 0.635*T_d_*.

Tables 1 and S1 compare the results from numerical integrations with this approximation; as expected from the assumptions made in deriving this approximation, it is most accurate when *γ* is large and *a* is not too close to 1. Overall, for low frequencies of recombination, *T_s_* is a non-negligible fraction of *T_d_*, but decreases towards zero with increasing rates of recombination, as would be expected.

**Table 1.**
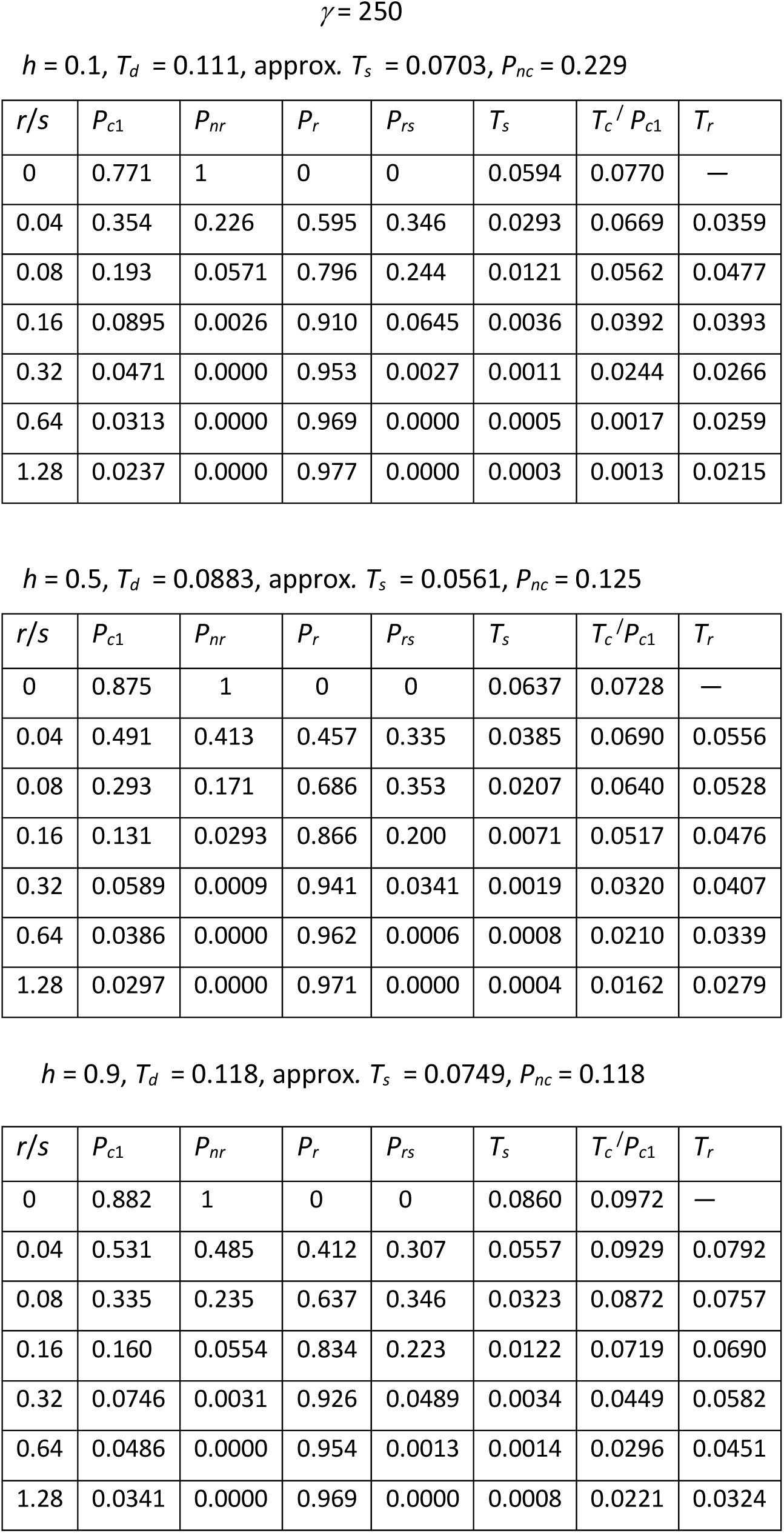
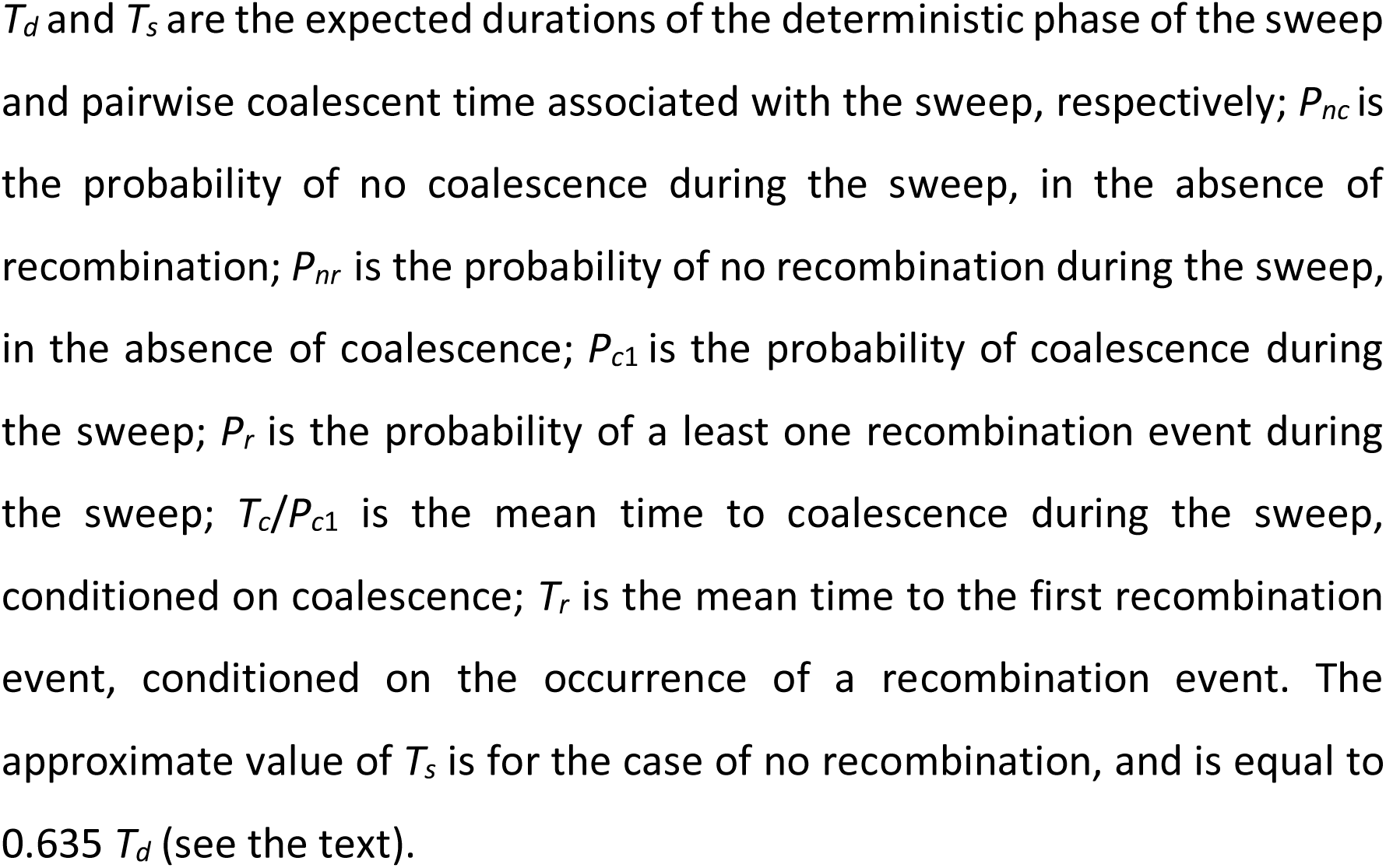
Parameters describing the effect of a single sweep.

#### Multiple recombination events

Finally, the problem of multiple recombination events needs to be considered. In principle, this problem can be dealt with on the lines of Equation 10, but this involves multiple integrals of increasing complexity as more and more possible events are considered. The following heuristic argument can be used instead. A first approximation is to assume that, if the frequency of recombination is sufficiently high, multiple recombination events are associated with a coalescent time equal to that of an unswept background, *B*_1_. In contrast, a single recombinant event is associated with a mean coalescent time of *B*_1_ +*T_d_*, since the recombinant cannot coalesce with the non-recombinant haplotype until the end of the sweep. If the probability of a single recombinant event is denoted by *P_rs_*, Equation 11a is replaced by:

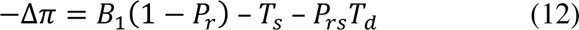

*P_rs_* is given by the probability of a recombination event that is followed by no further recombination events. This event requires both the recombinant A_1_ haplotype (whose rate of recombination at an A_2_ frequency of *x* is *ρx*) and the non-recombinant A_2_ haplotype (whose rate of recombination is *ρ*[1 – *x*]) to fail to recombine.

We thus have:

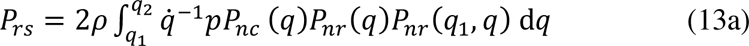

where *P_nr_*(*q*_1_, *q*), is the probability of no further recombination after an A_2_ frequency of *q*, given by:

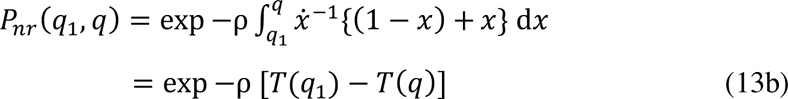

However, Equation 12 ignores the fact that there is a time-lag until the initial recombination event, whose expectation, conditioned on the occurrence of the initial recombination event, is denoted by *T_r_*. This lag contributes to the time to coalescence of multiple recombinant alleles, causing the reduction in diversity to be smaller than predicted by Equation 12b. The probability of multiple recombination events is (*P_r_* – *P_rs_*), so that a better approximation is to deduct (*P_r_* – *P_rs_*)*T_r_* from the left-hand side of Equation 12, giving:

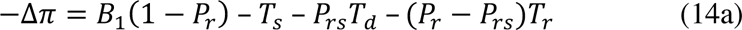

Where

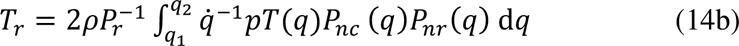

The integral for *T_r_* can be expressed in terms of *ρ*, *γ*, *a* and *b*, on the same lines as for Equation 10.

Equations 14 are likely to overestimate the effect of recombination on the sweep effect, as complete randomization of the sampled pair of haplotypes is unlikely to be achieved, whereas Equation 11a clearly underestimates it; Equation 12 should produce an intermediate prediction. The correct result should thus lie between the predictions of Equation 11 and Equation 14. When the ratio of the rate of recombination to the selection coefficient, *r*/*s*, is << 1, all three expressions agree, and predict a slightly smaller sweep effect than Equation 9 of HB.

### Comparisons with simulation results

Numerical results for Equations 11 can be obtained by numerical integration of the formulae given in the Appendix. For speed of computation, Simpson’s Rule with *n* + 1 points was used here; this method approximates the integral of a function by a weighted sum of discrete values of the integrand over *n* equally spaced subdivisions of the range of the function (Atkinson 1989). It was found that *n* = 200 usually gave values that were close to those for a more exact method of integration; for the results in the figures in this section, *n* = 2000 was used. Background selection effects are ignored here, so that *B*_1_ and *B*_2_ are set to 1. Simulation results for hard sweeps for an autosomal locus with random mating were obtained using the algorithm of Tajima (1990) (see the Methods section), providing a basis for comparison with the predictions based on Equations 11a and 14 (denoted by *C1* and *C2*, respectively), and on the star phylogeny approximation that ignores coalescence of non-recombined alleles during the sweep, Equation 11b (*NC*). The results are shown in Figure 1, with the reduction in diversity observed at the end of the sweep, –Δ*π*, on a log_10_ scale, plotted against *r*/*s* on a log_2_ scale, with values of *r*/*s* increasing by a factor of two from 0.04 to 1.28. Figure S1 of File S1 shows the same results on linear plots with *r*/*s* from 0 = 0.32, with the addition of the values of –Δ*π* for *r*/*s* = 0. Since *C1* and Equations 9 of HB, which ignore the term in *T_s_* in Equation 11a, gave very similar results, only the former are shown here.

One feature that is worth noting is that, with no recombination, the simulations and *C1/C2* formulae (which are identical and exact in this case) predict –Δ*π* values that are substantially less than one, especially with the lower strengths of selection. The *NC* approximation predicts a complete reduction in diversity, since the probability of coalescence is one and the duration of the sweep is ignored (this can be seen most clearly in the linear plots in Figure S1). In contrast, *NC* underestimates –Δ*π* when the recombination rate is high compared with the selection coefficient; even for *r*/*s* as small as 0.16 there can be a very large ratio of the simulation value to the *NC* value, although the simulation value of –Δ*π* is then usually quite small (10% or less) for this value of *r*/*s*. For example, with *γ* = 250, *h* = 0.5 and *r*/*s* = 0.16, the simulation value of –Δ*π* was 0.0959 (s.e. 0.0019), whereas the *NC* value was 0.0293. Conversely, *C1* tends to overestimate Δ*π* for the higher values of *r*/*s*, reflecting the fact that it does not allow for multiple recombination events.

Overall, *C2* agrees quite well with most of the simulation results, especially for *h* = 0.5, but tends to underestimate –Δ*π* for *h* = 0.9, especially for large *r*/*s*, presumably because the relatively long period which A_2_ spends at high frequencies means that a substantial proportion of multiple recombination events involve a return of a recombined neutral site back onto the A_2_ background, For much larger *r*/*s* values than are shown here, *C2* can become negative, indicating that it over-corrects for multiple recombination events, but –Δ*π* is then very small, so the effect is probably not biologically important. As has been found previously (Teshima and Przeworski 2006; Ewing *et al*. 2011; Hartfield and Bataillon 2020), –Δ*π* increases with *h* for low values of *r*/*s*, but the values for each *h* converge as *r*/*s* increases.

Tables 1 and S1 show details of some of the relevant variables, obtained by numerical integration. They confirm the conclusion that there can be a substantial probability of coalescence during the sweep, as given by *P_c_*_1_ in Equation 8a; this probability decreases much more slowly with *r*/*s* than does the probability of no recombination in the absence of coalescence (*P_nr_*). In parallel with this behavior of *P_c_*_1_, the unconditional probability of no recombination, 1 – *P_r_*, decreases much more slowly with *r*/*s* than *P_nr_*. This explains why the *NC* approximation for the reduction in diversity performs rather poorly at high *r*/*s* values.

The results also show that the probability of a single recombination event (*P_rs_*, given by Equation 13a) becomes very small compared with the probability of at least one recombination event (*P_r_*, given by Equation 10a) as *r*/*s* increases, so that neglecting the effects of multiple recombination events leads to errors in predicting sweep effects on diversity. For high values of *r*/*s*, the conditional mean times to coalescence and to the first recombination event are both small relative to the duration of the sweep, implying that these events must occur quite soon if they are to occur at all.

To illustrate the approximations further, both Tajima and HB simulation values of – Δ*π* for an autosomal locus in a randomly mating population with three difference dominance coefficients and *γ* = 500, together with the theoretical predictions, are shown in Figure S2. These confirm the general conclusions from Figure 1, despite the fact that the HB simulation results seem to be considerably noisier than the Tajima results, sometimes showing a non-monotonic relation between –Δ*π* and the recombination rate.

Figure 2 displays results for selfing rates of 0.5 and 0.95, corresponding to *F* values of 0.3333 and 0.9048, respectively. The reduction in diversity is plotted against the scaled recombination rate for an autosomal locus with outbreeding; the scaled effective recombination rate with inbreeding is much smaller than this, as described above. Here, only the HB simulation results are shown, as the Tajima method cannot give an exact representation of the system with non-random mating. As before, *C1* and the approximation given by Equations 9 of HB mostly give very similar predictions, whereas *C2* predicts smaller effects that agree slightly less well with the simulations at the higher recombination rates. However, for *S* = 0.95, especially with *h* = 0.5, the simulations yield considerably larger sweep effects at relatively high recombination rates than any of the theoretical predictions. This presumably reflects the fact that random variation among individuals in the occurrence of selfing versus outcrossing events means that individuals sampled in a given generation differ in the numbers of generations of selfing in their ancestral lineages, and hence in the extent to which recombination and selection have interacted to cause departures from neutral expectations (Kelly 2007). This is not taken into account in the formula used to correct for the effects of selfing on the effective rate of recombination (Equation 1 in HB).

**Figure 2.**
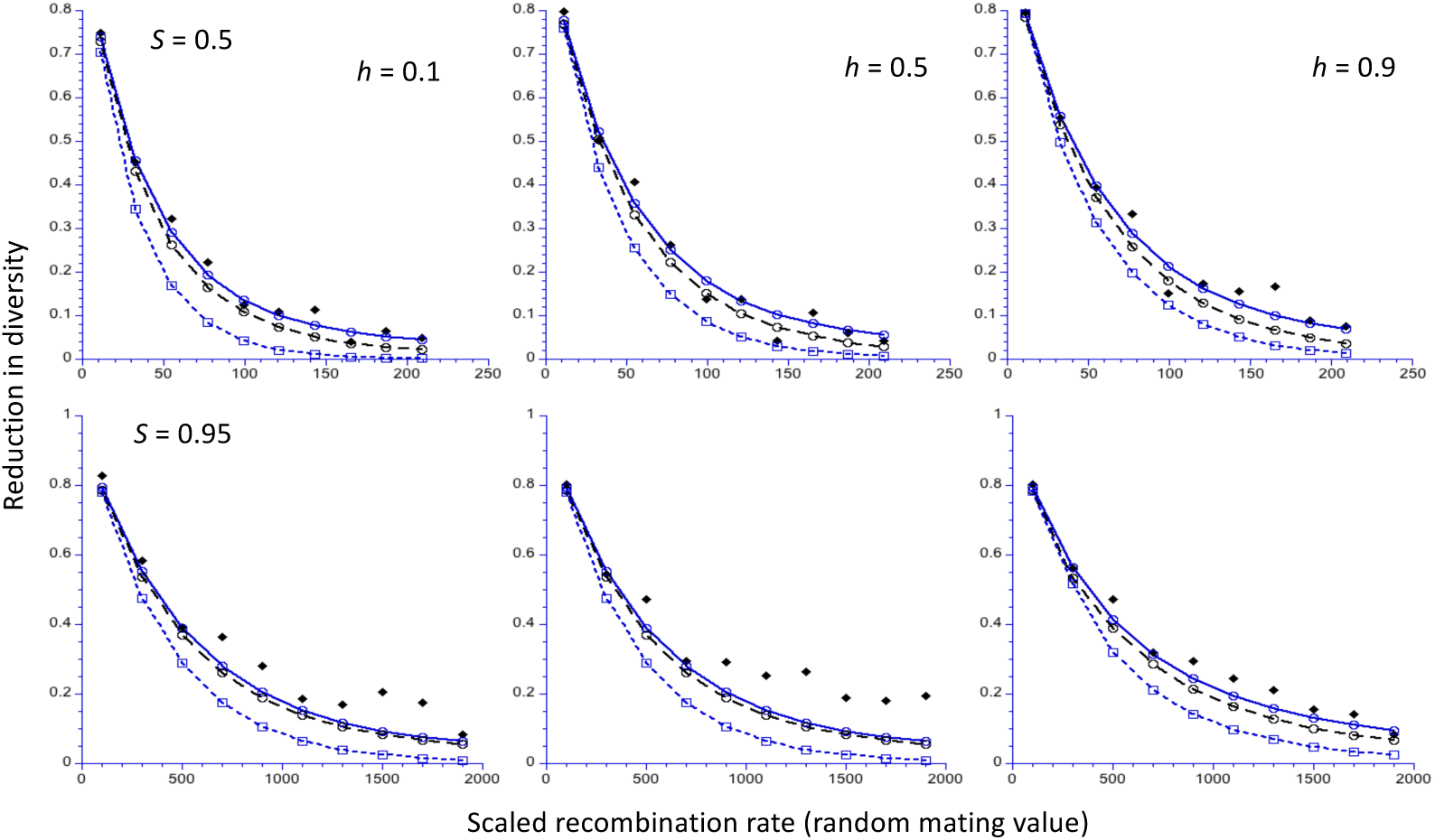
The reduction in diversity (relative to the neutral value) at the end of a sweep for an autosomal locus, as a function of the scaled rate of recombination (2*N_e_r*) for a randomly mating population. The results for two different selfing rates, *S*, (upper and lower panels) are shown, together with three different values of the dominance coefficient, *h*. A population size of 5000 is assumed, with a scaled selection coefficient for A_2_A_2_ homozygotes in a randomly mating population (*γ* = 2*N_e_s*) of 500. The filled black lozenges are the mean values from computer simulations (Hartfield and Bataillon 2020); the open blue circles and open black circles are the *C1* and *C2* predictions, respectively; the open blue squares are the *NC* predictions.

#### Inaccuracy of the NC approximation

Given the widespread use of the star phylogeny assumption in methods for detecting recent sweeps and inferring the parameters of positive selection, described in the introduction, it is disconcerting that the *NC* approximation is systematically somewhat inaccurate with respect to pairwise diversity at relatively large values of *r*/*s*. Some insights into this effect can be obtained from examining an approximation for the case of autosomal inheritance with *h* = 0.5 and random mating, which is derived in the Appendix.

If we write *R* = 4*r*/*s* and *α* = 2/*γ*, and assume that *R* > 1 and *γ* >> 1, the reduction in diversity after a sweep is close to one minus the probability of recombination during the sweep, as given by Equation 11a with *T_s_* = 0. From Equation A7, we have:

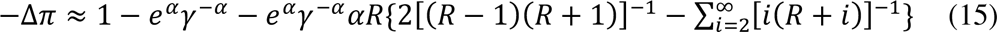

This series converges quite slowly when *R* is large, but the formula agrees well with the results of numerical integration even for *r*/*s* as low as 0.4, provided that *γ* is sufficiently large. It tends to break down for high values of *R* (> 10), especially for relatively small *γ*. Equation 15 implies that, paradoxically, *larger* values of *γ* lead to *smaller* values of the diversity reduction for a given *r*/*s*, as can be seen as follows. For large *γ*, 1 – *e^α^γ* ^−^*^α^* ≈ *α* ln(*γ*) for large *γ*, which is nearly proportional to *α*. In addition, the term in braces is positive for sufficiently large *R*, and its product with *αe^α^γ* ^−^*^α^* is also nearly proportional to *α*.

This effect can be seen in Figures 1 and S1. A doubling of *γ* results in substantially smaller values of the diversity reduction for a given value of *r*/*s*, as expected from the above properties of Equation 15. This applies to both the *C1* and *C2* predictions, as well as the simulations. Thus, contrary what is predicted by the *NC* approximation, the effect of a sweep on diversity is negatively related to the scaled strength of selection, for a given value of *r*/*s*. The intuitive interpretation of this finding is that weaker selection prolongs the duration of a sweep, allowing more opportunity for coalescence versus recombination.

## The effects of recurrent selective sweeps on nucleotide site diversity

### Theoretical results

The approach of CC for determining the effects of recurrent sweeps, which was based on the *NC* approximation, can be modified to apply to the more general case considered here. It is assumed that adaptive substitutions occur at a total rate *ω* per 2*N_e_* generations, such that the times between substitutions follow an exponential distribution with rate parameter *ω* (this rate includes any effects of BGS in reducing the probability of fixation of favorable mutations). By summing up over all relevant nucleotide sites that contribute to the effect of sweeps at a focal neutral site, weighting each selected site by its rate of adaptive substitution (which may differ according to the class of site subject to selection), and then dividing by *ω*, we can define expected values of *P_r_*, *P_r s_*, *T_s_*, *T_r_* and *T_d_* for a given neutral site (expected values are denoted by overbars in what follows).

As a first step, it is useful to note that Equation 8 of CC for the expected nucleotide site diversity immediately after a substitution, *π*_0_, is equivalent to:

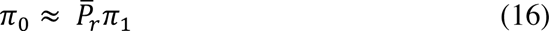

where *π*_1_ is the expected nucleotide site diversity at the time of the initiation of a new substitution. Both *π*_0_ and *π*_1_ are measured relative to the neutral diversity *θ*, and hence are equivalent to mean pairwise coalescent times relative to the neutral value, 2*N_e_*. This expression assumes that there is at most a single recombination event, and that a pair of alleles that have been separated by recombination onto the A_1_ and A_2_ backgrounds have a coalescent time equivalent to that for a pair of alleles that are sampled at the start of the sweep.

If we apply the argument leading to Equation 11a to take into account the lag time to coalescence of a pair of alleles separated by recombination, we obtain the *C1* approximation:

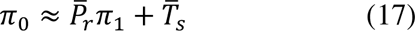

If we use the approach of Equations 14 for modeling multiple recombination events, we obtain the *C2* approximation:

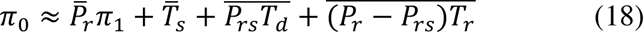

A somewhat more accurate expression can be found by noting that, under the assumption that multiple recombination events cause randomization between A_1_ and A_2_ haplotypes, so that coalescence occurs at rate *B*_1_^−1^, diversity will increase from its value at the start of a sweep over a time interval that is approximately the same as the difference between the sweep duration and the time of the first recombination event (see Equation 8b of the Appendix); this overestimates the time available for the increase in diversity, since a time greater than*T_r_* is required for randomization to occur.

These expressions enable us to find the expected diversity, *π*, for a pair of alleles sampled at a random point in time. This is done by assuming that such a time point falls between two sweeps, and that the length of the interval *T* separating the two sweeps follows an exponential distribution with parameter *ω*. Conditional on *T*, the time *τ* from a random sample to the first of the two sweeps is a uniform variate on the interval *T,* with p.d.f. equal to 1/*T*. The expected diversity at time *τ* (on the coalescent timescale) is given by the equivalent of Equation 9 in CC:

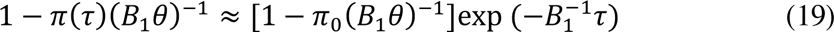

The overall expected diversity is thus given by:

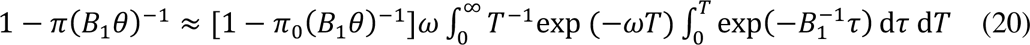

This expression is identical with Equation 10 of CC, so that their Equations 12 for *π* can be used, but with a more precise interpretation of the meaning of *π*.

### Comparisons with simulation results

The accuracy of Equations A10-A13 was tested using the simulation results from Figure 4 and Table S6 in CC. These simulations modeled a group of 70 linked genes with properties similar to those of typical *D. melanogaster* autosomal genes, and provided values of the mean nucleotide site diversity at synonymous sites under the assumption that they are selectively neutral. The genetic model and parameters of the simulations are summarized in the Methods section of this paper; full details are given in CC. The effect of BGS on the rate of substitutions of favourable mutations for a given parameter set was calculated here by multiplying the rate in the absence of sweeps by the value of *B*_1_ for neutral sites obtained from simulations; use of *B*_2_ instead of *B*_1_ made little difference. The corresponding theoretical predictions were obtained for a single gene with the structure described in the Methods section, on the assumption that sweep effects decay sufficiently fast with distance from the selected site that each gene can be treated independently; this is probably not entirely accurate for the lowest rate of crossing over studied here.

Equations A10-13 can be applied in two ways. First, the mean values of the relevant quantities across all neutral sites can be determined, and substituted into these equations, the procedure used in CC for the older method of predicting recurrent sweep effects. Second, the values of these statistics for individual neutral sites can be used to predict *π*, and the mean of *π* taken across all neutral sites in a gene, as described in the Methods section. The latter procedure is more accurate statistically, and is used here for the *C2* predictions; in practice, the two methods yield similar results for the parameter sets used here.

Figure 3 shows the reductions in neutral diversity per gene obtained from the simulation results (red bars), the *C2* predictions using Equations A10-A13 (blue bars), the predictions from Equations 12 of CC (black bars) that use the *NC* approximation, and the *NC*-based formula for recurrent sweep effects that assumes competing exponential processes of coalescence due to drift and selection (Equation 7 of CC) (white bars). For the last two calculations, the *NC* assumption with the deterministic sweep duration (*T_d_*) was used to calculate the effect of a sweep, using the first method described above for estimating the mean effect over neutral sites. Further details of the calculations are given in the Methods section.

**Figure 3.**
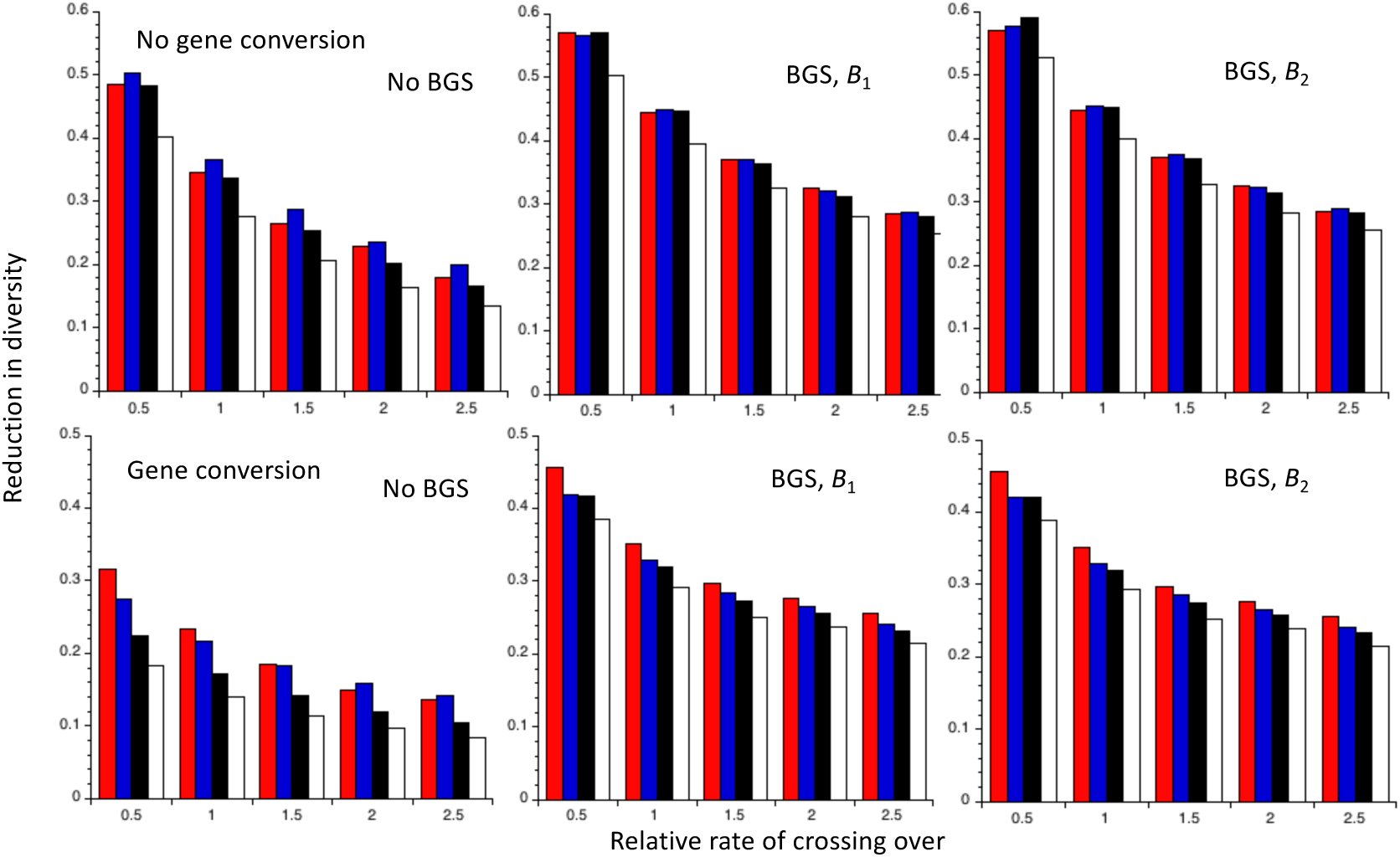
Comparisons with several different theoretical predictions of the mean values of the reduction in diversity obtained in computer simulations of recurrent sweeps with random mating, autosomal inheritance and semi-dominant favorable mutations, described in CC. The X axis shows the values of the rate of crossing over, expressed relative to the mean value for *Drosophila melanogaster.* The red bars are the mean values of the simulation results for neutral (synonymous sites) in a group of 70 genes; the blue bars are the predictions from Equations A10-13; the black bars are the predictions from Equation 12b of CC; the white bars are the predictions from the standard coalescent model of recurrent sweeps, assuming no coalescence within sweeps (Equation 7 of CC). Cases with and without gene conversion (upper and lower panels, respectively), and in the absence or presence of background selection (BGS), using either *B*_1_ or *B*_2_ to predict fixation probabilities when BGS is acting, are shown.

The most notable point is that, as was also found by CC, the last method is consistently the least accurate, especially at low CO rates in the presence of gene conversion but without BGS. In the absence of gene conversion, the predictions from Equation 12 of CC and Equations A10-13 generally have a similar level of accuracy. However, in the presence of gene conversion, the latter predictions perform the best, and provide fairly accurate predictions except for the lowest CO rate. This comparative failure of predictions based on the *NC* approach presumably reflects the fact that it considerably underestimates the effects of sweeps in the presence of recombination, as was seen in Figure 1. Given that gene conversion is pervasive in genomes, and contributes substantially to recombination rates over short physical distances, the approach developed here should provide the most accurate predictions of recurrent sweep effects, despite its heuristic basis. The inaccuracy of the predictions based on the competing exponential process model, introduced by Kaplan *et al*. (1989) and further analyzed by Stephan *et al*. (1992), Kim and Stephan (2000) and Gillespie (2002), is probably due to the fact that it treats sweeps as point events, allowing too much opportunity for drift-induced coalescent events between sweeps (CC, p.293).

## X chromosomes versus autosomes

### Introduction

There has been considerable interest in comparing the properties of variability of sequences on X or Z chromosomes with those on autosomes (A), since these may shed light on questions such as the relative importance of BGS versus selective sweeps in shaping genome-wide patterns of variability, and on the causes of the apparently faster rates of adaptive evolution on the X or Z chromosome (Charlesworth *et al*. 2018; Wilson Sayres 2018). It therefore seems worth revisiting this question in the light of the models of selective sweeps developed here, which can easily be applied to sex-linked loci. The findings extend those of Betancourt *et al*. (2004), who considered only the case of selection acting equally on the two sexes and used the equivalent of the *NC* model described above.

As noted by Betancourt *et al*. (2004), there are important differences in the theoretical expectations for taxa such as *Drosophila* and Lepidoptera, in which autosomal recombinational exchange is absent in the heterogametic sex, and taxa such as mammals and birds, where recombination is absent between the X (Z) and Y (W), but occurs on autosomes and pseudoautosomal regions in the heterogametic sex. In the first type of system, the sex-averaged effective rate of recombination (which controls the rate of breakdown of linkage disequilibrium) between a pair of X-or Z-linked genes is 4/3 times that for an autosomal pair with the same rate of recombination in the homogametic sex, due to the fact that the X or Z spends 2/3 of its time in the homogametic sex and 1/3 of its time in the heterogametic sex, whereas an autosome spends half of its time in the heterogametic sex where it cannot recombine (Langley *et al*. 1988). In the second type of system, the ratio of sex-averaged rates is 2/3 (Betancourt *et al*. 2004). These two systems will be referred to here as the “*Drosophila*” and “mammalian” models, respectively. For brevity, only male heterogamety is considered here; the results for female heterogamety can be obtained by interchanging male and female.

### The effect of a single sweep on X-linked diversity

It is straightforward to use the framework leading to Equation 9 to examine the effect of a single sweep on variability for an X-linked locus. In this case, it is necessary to model the effects of sex differences in the effects of a mutation on male and female fitnesses, since these greatly affect the evolutionary trajectories of favorable X-linked mutations (Rice 1984; Charlesworth *et al*. 1987; Charlesworth 2020). Three extreme cases are considered here: no sex-limitation of fitness effects (so that the homozygous selection coefficient, *s*, is the same for males and females), male-only fitness effects, and female-only fitness effects. Random mating is assumed throughout.

For simplicity, the dominance coefficient *h* is assumed to be independent of sex. For the autosomal case with weak selection, a single *s* that is given by the mean of the male and female fitness effects is sufficient to describe the system (Nagylaki 1979). The values of the coefficients *a* and *b* in Equation 6 for the three types of sex-dependent fitness effects can be obtained from the expressions in Box 1 of Charlesworth (2020). With no sex-limitation and random mating, *a* = (2*h* + 1)/3 and *b* = 2(1– 2*h*)/3; with male-only selection, *a* = 1/3 and *b* = 0; with female-only selection, *a* = 2*h* /3 and *b* = 2(1 – 2*h*)/3. In order to ensure comparable strengths of selection for X and A with the same patterns of relation between gender and fitness, the values of *s* for the cases of male- and female-only effects with X-linkage are set equal to twice the corresponding autosomal *s* without sex-limitation, compensating for the fact that the effective *s* for a sex-limited autosomal mutation is only one-half of the selection coefficient in the affected sex.

Figure 4 shows the reductions in diversity at the end of a sweep predicted by the *C1* and *C2* methods for the *Drosophila* model, together with the results of simulations using the algorithm of Tajima (1990), for the case of an X-linked locus whose effective population size, *kN_e_*_0_, is three-quarters of that for A, *N_e_*_0_. This case corresponds to a randomly mating population in which males and females have equal variances in reproductive success (Wright 1931). With *h* = 0.5 and *k* = 0.75, all three types of sex-specific selection on X-linked loci have similar evolutionary dynamics, provided that the selection coefficients are adjusted as described above (Charlesworth 2020). No differences among their sweep effects are thus to be expected, apart from small deviations reflecting numerical inccuracies in the integrations. This expectation is confirmed by the results shown in Figure 5. As before, the *C1* approximation predicts much larger effects than *C2* at high *r*/*s* values; the *NC* approximation predicts even smaller effects than *C2* (results not shown).

**Figure 4.**
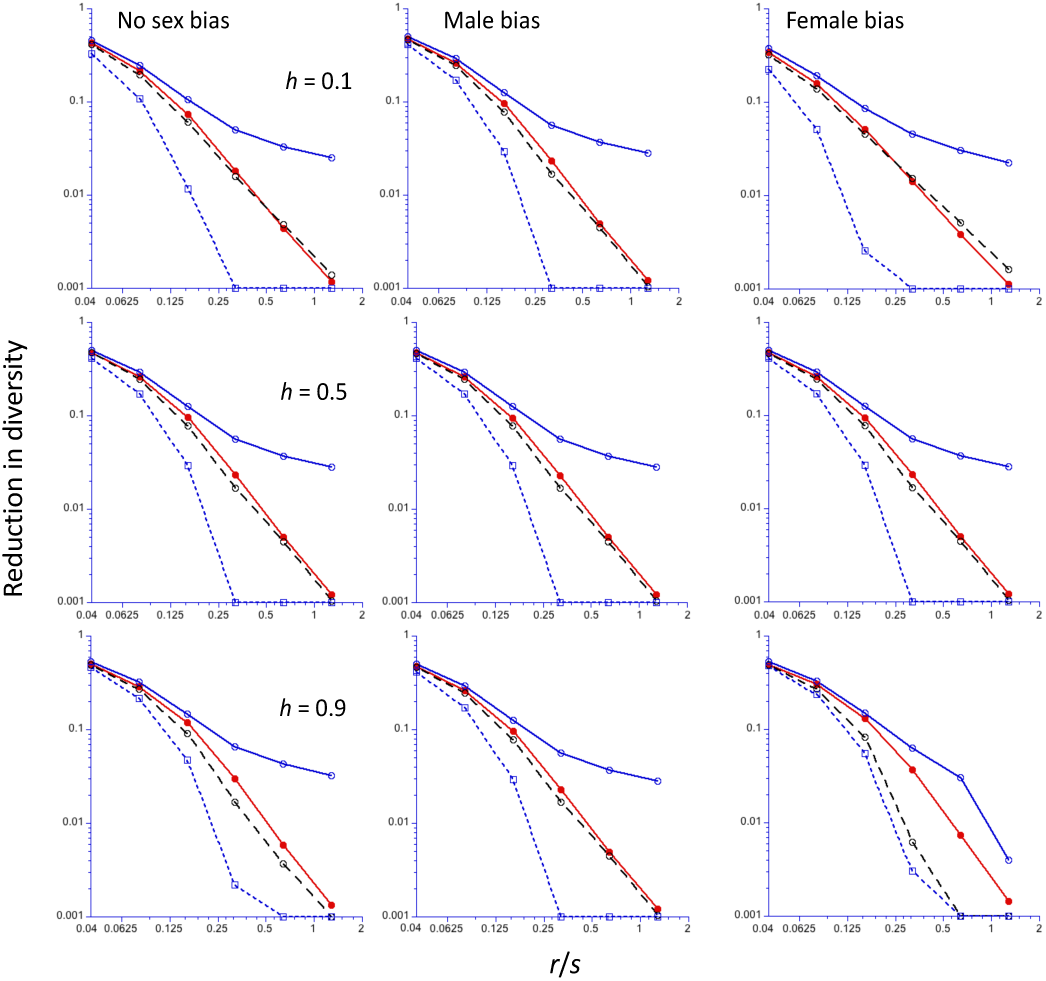
The reduction in diversity (relative to the neutral value) at the end of a sweep for an X-linked locus (Y-axis, log_10_ scale), as a function of the ratio of the frequency of recombination (*r*) to the selection coefficient for homozygotes (*s*) (X-axis, log_2_ scale). The *Drosophila* recombination model is assumed; *N_e_* for the X chromosome is three-quarters of that for the autosomes. The results for mutations with no sex limitation are shown in the left-hand panels; those for male-limited and female-limited mutations are shown in the middle and right-hand panels, respectively. A population size of 5000 is assumed, with a scaled selection coefficient for an autosomal mutation in a randomly mating population (*γ* = 2*N_e_s*) of 250 for the cases with no sex-limitation. For the sex-limited cases, *γ* = 500 to ensure comparability to sex-limited autosomal mutations. Results for three different values of the dominance coefficient (*h*) are shown, with *h* increasing from the top to bottom panels. The filled red circles are the mean values from computer simulations, using the algorithm of Tajima (1990); the filled blue circles and black circles are the *C1* and *C2* predictions, respectively; the open blue squares are the *NC* predictions. Values of the reduction in diversity less than 0.001 have been reset to 0.001.

**Figure 5.**
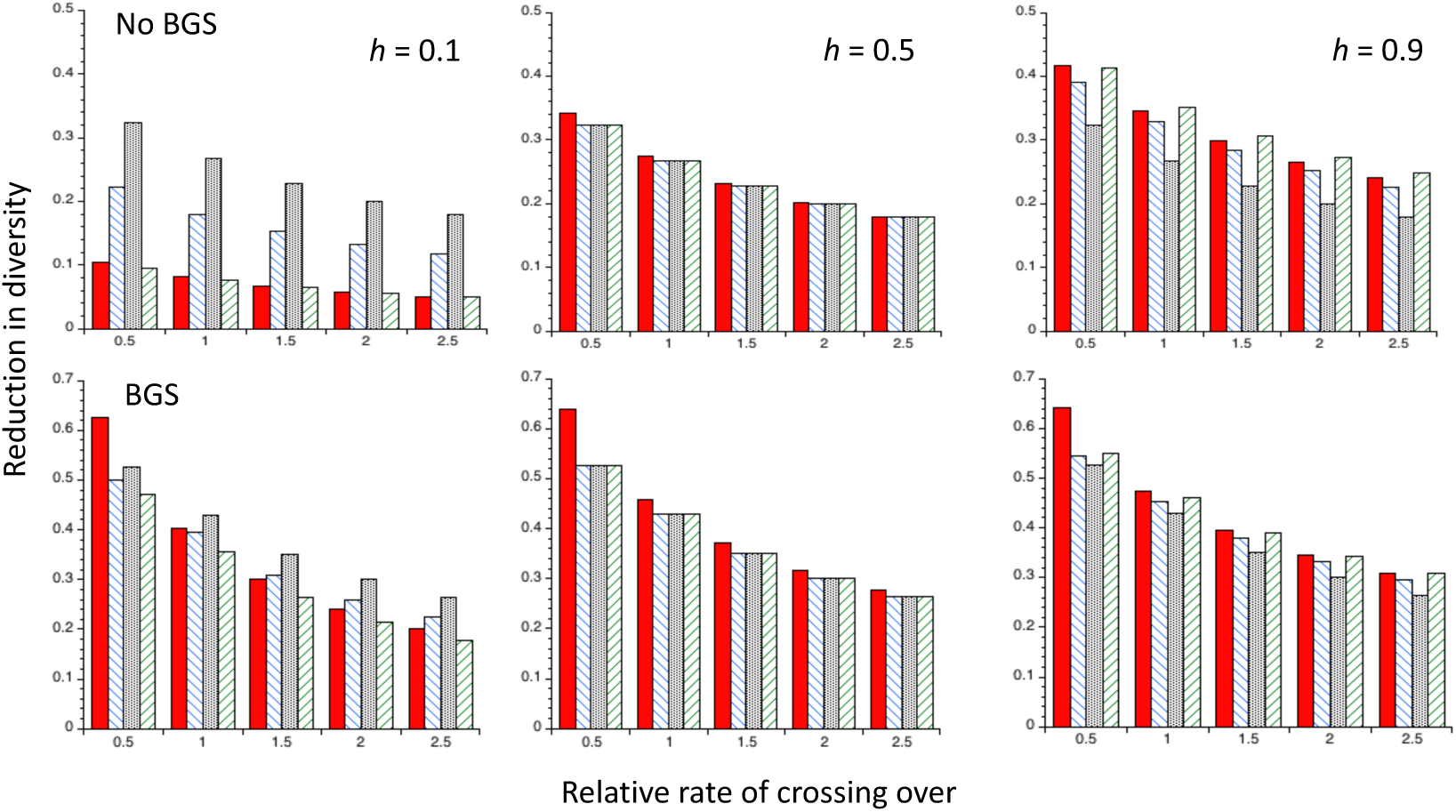
Reductions in diversity (relative to neutrality) under recurrent sweeps at autosomal and X-linked loci for the *Drosophila* model, using the *C2* theoretical predictions with gene conversion and five different rates of crossing over relative to the autosomal standard value (the X-linked rates of crossing over and gene conversion were chosen to give the same sex-averaged effective rates as for the autosomes). *N_e_* for the X chromosome is three-quarters of that for the autosomes. The upper panel is for cases without BGS; the lower panel is for cases with BGS (using the parameters described in the main text). The filled red bars are for autosomal mutations, the hatched blue bars are for X-linked mutations with no sex-limitation, the stippled black bars are for male-limited X-linked mutations, and the hatched green bars are for female-limited X-linked mutations. For the sex-limited cases, *γ* = 500 to ensure comparability with the autosomal and non-sex-limited X-linked mutations.

The comparison of the X-linked results with *h* = 0.5 with the autosomal results for *γ* = 250 in Figure 1 confirms the expectation that the diversity reductions are the same for the two genetic systems, when *s* is adjusted appropriately. In addition, female-limited X-linked mutations have the same effects as female-limited autosomal mutations for all *h* values, again as expected from their similar dynamics. For a given *r*/*s*, male-limited selection gives the largest reduction in X-linked diversity when *h* = 0.1, which is substantially larger than the autosomal and female-limited values for the same adjusted selection strength. This is expected from the slow initial rates of increase in the frequencies of partially recessive autosomal or female-limited X-linked mutations (Haldane 1924; Van Herwaarden and van der Wal 2002; Teshima and Przeworski 2006; Ewing *et al*. 2011; Charlesworth 2020). With *h* = 0.9, the differences between the various cases are relatively small, with male-limited X-linked mutations having the smallest effects, and non-sex-limited and female-limited mutations having almost identical effects. The sweep effects decrease more rapidly with *r*/*s* for X than for A, as expected from the higher effective recombination rate for X.

Figure S3 shows comparable results for the mammalian model. The results are broadly similar to those for the *Drosophila* model, the main difference being that the sweep effects are always larger than for the corresponding *Drosophila* cases, as would be expected from the fact that the effective rate of recombination on the X chromosome is half the *Drosophila* value. In this case, the X sweep effects decrease more slowly with *r*/*s* than the A effects. Figure S4 in File S1 shows the results for both the *Drosophila* and mammalian models on a linear scale.

The X/A ratio of *N_e_* values in the absence of selection at linked sites (*k*) may differ from three-quarters. Sex differences in these variances cause *k* values that differ from 0.75 (Caballero 1995; Charlesworth 2001; Vicoso and Charlesworth 2009), with higher male than female variances leading to *k* > 0.75, and lower male than female variances having the opposite effect. Male-male competition for mates is likely to cause a higher male than female variance in fitness, so that *k* can be greater than 0.75 with male heterogamety. In contrast, female heterogamety with sexual selection leads to *k* < 0.75. Some examples of the reductions in diversity at the end of a sweep with *k* ≠ 0.75 using the *C2* predictions are shown in Figure S5 in File S1, for the *Drosophila* and mammalian models, respectively. Comparing these with Figures 1 and S1, it can be seen that smaller *k* values cause somewhat larger X-linked sweep effects for all modes of selection and for both genetic systems. The effects are, however, relatively small, and unlikely to be detectable in most datasets. This pattern is presumably caused by the fact that smaller *k* means that coalescence during a sweep occurs more rapidly relative to recombination.

### The effects of recurrent selective sweeps on X-linked diversity

Expressions for the effects of recurrent selective sweeps on X-linked neutral diversity can be obtained using the appropriate modifications to Equation 17 (the *C1* approximation) and Equations A11-A13 (the *C2* approximation). There is an important factor that leads to differences between A and X sweep effects, additional to those considered in the previous section. This is the fact that the expected coalescent time for X is 2*kN_e_*_0_ instead of 2*N_e_*_0_, with *k* generally expected to be less than 1 (Charlesworth 2001; Vicoso and Charlesworth 2009). The parameter *ω* that appears in the equations for sweep effects is the expected number of substitutions over 2*kN_e_*_0_ generations, so that a smaller value of *k* for a given rate of substitution per generation implies a smaller *ω* value. An alternative way of looking at this effect is to note that *k* < 1 implies a faster rate of genetic drift for X than A; other things being equal, coalescent events induced by drift are then more frequent for X than for A, relative to coalescent events induced by selection (Betancourt *et al*. 2004).

A countervailing factor is that the rates of substitution per generation of favorable X-linked mutations are expected to be higher than for comparable autosomal mutations with male-limited or non sex-limited selection, given sufficiently small *h* values (the condition is *h* < 0.5 with *k* = 0.75) (Charlesworth *et al*. 1987, 2018; Vicoso and Charlesworth 2009); expressions for the rate of substitution are given in the final section of the Appendix. These opposing effects of sex linkage implies that simple generalizations about the effects of sweeps on X-linked versus autosomal variability cannot easily be made, as will shortly be seen.

Figure 5 shows the *C2* predictions for the reductions in diversity relative to neutral expectation for X-linked and autosomal loci under the *Drosophila* model, as a function of the ratio of the autosomal effective CO rate to a value of 2 x 10^−8^ per basepair for *D. melanogaster*, using the same gene structure that was used to generate the theoretical results for autosomes shown in Figure 3 (see Methods section). Gene conversion was allowed at the same rate of initiation as crossovers. The CO and gene conversion rates for the X chromosome were set by multiplying the corresponding autosomal effective rates by 0.75, so that the effective recombination rates for the X chromosome and A are equal, following the procedure used in empirical comparisons of diversity levels on the X and A (Campos *et al*. 2014). Here *k* = 0.75, so that potential effects of sexual selection or variance in female reproductive status are absent.

As described in the Methods section, the values of the BGS parameter *B*_1_ were obtained from estimates given by Charlesworth (2012), which include contributions from selectively constrained non-coding sequences as well as coding sequences; comparable values were obtained in the more detailed analyses of Comeron (2014). As described in CC, the BGS effect parameter *B*_1_ for the X chromosome with a relative crossing rate of 0.5 was set to a relatively high value (0.549 instead of 0.449) to correct for the relatively low gene density in this regions of the *D. melanogaster* X chromosome, whereas the values for the rates of 1, 1.5, 2 and 2.5 assumed normal gene densities, giving *B*_1_ values of 0.670, 0.766, 0.818 and 0.852, respectively. This results in a relatively weak effect of selection in reducing diversity for the lowest CO rate compared with the autosomes, where the *B*_1_ values were set to 0.538, 0.733, 0.813, 0.856 and 0.883 for the relative CO rates of 0.5, 1, 1.5, 2 and 2.5, respectively. Male and female mutation rates were assumed to be equal, in view of the lack of strong evidence for a sex difference in mutation rates in *Drosophila* (Charlesworth *et al*. 2018). For convenience, *B*_2_ was assumed to be equal to *B*_1_.

As expected, in the absence of BGS the X results for *h* = 0.5 (upper panel of Figure 5) are the same for the three types of sex-specific fitness effects. The X effects are slightly smaller than the A effects for low recombination rates, reflecting the reduced rate of substitution on the coalescent timescale of the lower *N_e_* for X than A, which was described above. With *h* = 0.1, the lower rates of substitution of A mutations and female-limited X mutations greatly reduce their sweep effects, but the effects for non-sex-limited and male-limited X mutations are much larger than for A mutations. With *h* = 0.9, male-limited X mutations have the weakest effects, while the other three classes of mutations have quite similar effects, with female-limited mutations having the largest effects. Similar general patterns are seen with BGS, with much smaller differences between X and A than in the absence of BGS, except for the lowest CO rate, where the relatively small BGS effect for the X causes it to have a much smaller reduction in diversity compared with A. Comparable results with no gene conversion are shown in Figure S6 of File S1. The general patterns are quite similar to those with gene conversion, but with a greater sensitivity to the CO rate with *h* = 0.1 and no sex-limitation or male-limitation, especially with no BGS.

Figure 6 shows the values of the X/A ratio of diversities (*R_XA_*) for different CO rates, obtained from the results shown in Figure 5; Figure S7 shows comparable results with no gene conversion. First, consider the case when BGS effects are absent. With *h* = 0.5, *R_XA_* is always close to the neutral expectation of 0.75 for all modes of selection; this is also true with female-limited selection for all three dominance coefficients, with a slight tendency towards *R_XA_* > 0.75 for low CO rates, declining towards 0.75 as the CO rate increases. With *h* = 0.1 and no sex-limitation, it can be seen that *R_XA_* << 0.75 for the lowest CO rates, approaching 0.7 at the highest rate. With *h* = 0.1 and male-limitation, *R_XA_* increases with the CO rate, but remains well below 0.7 even at the highest rate. With *h* = 0.9 and no sex-limitation, *R_XA_* is slightly larger than 0.75 for the lowest rate of crossing, approaching 0.75 as the rate increases; with male-limited selection, *R_XA_* > 0.85 at the lowest CO rate, and *R_XA_* ≈ 0.8 at the highest rate.

**Figure 6.**
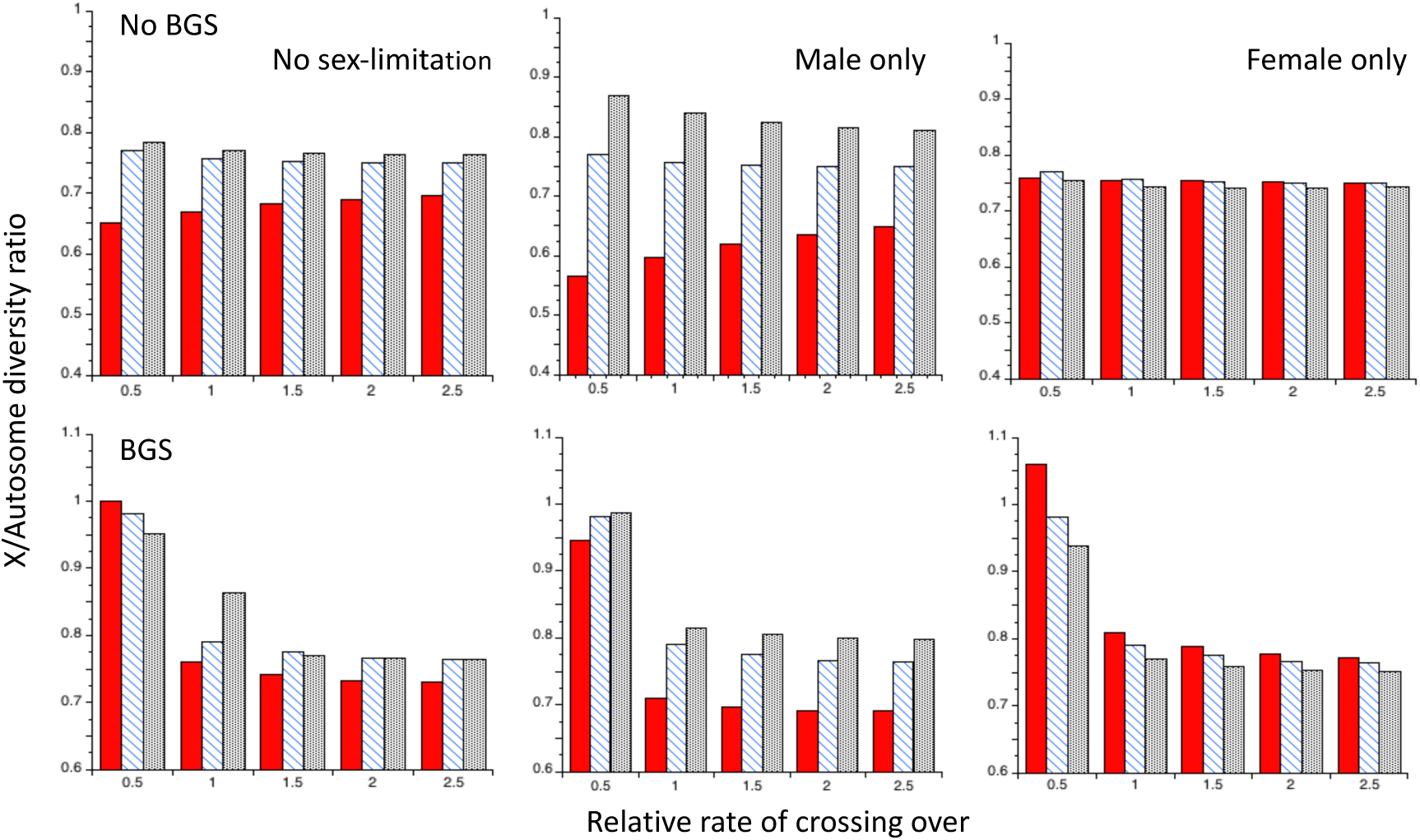
The ratios of X chromosome to autosome nucleotide site diversities (*R_XA_*) for the *Drosophila* model under recurrent sweeps, using the *C2* theoretical predictions with gene conversion and five different rates of crossing over relative to the autosomal standard value (the X-linked rates of crossing over and gene conversion were chosen to give the same sex-averaged effective rates as for the autosomes). *N_e_* for the X chromosome is three-quarters of that for the autosomes (*k =* 0.75). The upper panel is for cases without BGS; the lower panel is for cases with BGS. The filled red bars are for *h* = 0.1 (the dominance coefficient of favorable mutations), the hatched blue bars are for *h* = 0.5 and the black stippled bars are for *h* = 0.9. The other details are as for Figure 5.

The presence of BGS greatly alters these patterns; the lower gene density for the X in the region with the lowest CO rate causes *R_XA_* values of 0.9 or more for all three modes of selection and dominance coefficients. *R_XA_* even exceeds 1 for female-limited selection with *h* = 0.1 and the lowest CO. BGS causes a much steeper decline in *R_XA_* with the CO than in its absence (when it can even increase), especially with *h* = 0.1 and female-limited selection. The contrast between the presence and absence of BGS and the male-limited and non-sex-limited cases with *h* = 0.1 is especially striking. However, if the *B*_1_ value of 0.449 for a normal gene density is used for the lowest CO rate, X-linked diversity is considerably increased and *R_XA_* is correspondingly reduced; for example, with *h* = 0.1 and no sex-limitation, *R_XA_* = 0.849 instead of 1.001. Again, the patterns are similar to those found with gene conversion; with *h* = 0.1 and no sex-limitation or male-limitation, *R_XA_* is much more sensitive to the CO rate than when gene conversion is acting.

The effects of differences in *k* are shown in Figures S8 and S9, for the case with both BGS and gene conversion. Under the substitution model used here, a larger *k* is associated with a faster rate of substitution, countering the small effect of the size of a individual sweep described above. Comparisons with Figure 6 show that there tend to be somewhat larger effects of X-linked sweeps with the larger values of *k*. These translate into noticeably larger values of the X/A diversity ratios, but a reduced sensitivity of these ratios to the CO rate.

## Discussion

### General considerations

As described in the introduction, a widely-used simplification for calculating the effect of a selective sweep on nucleotide site diversity at a linked neutral site is the “star-like phylogeny” assumption that alleles sampled at the end of a sweep, and which have not recombined onto a wild-type background, coalesce instantaneously. Their mean coalescent time (relative to the purely neutral value) for a pair of alleles can then be equated to the probability that one of them undergoes a recombination event that transfers the neutral site onto the wild-type background (Wiehe and Stephan 1993; Barton 1998, 2000; Durrett and Schweinsberg 2004).

The results presented here show that this often leads to inaccuracies in predictions concerning the mean coalescent time for a pair of swept alleles, especially when the ratio of recombination rate to the homozygous selection coefficient (*r*/*s*) is relatively high, consistent with previous findings (Barton 1998; Hartfield and Bataillon 2020), as can be seen in Figures 1 and S1. Similarly, Figure 3 shows that with recurrent sweeps, gene conversion and no BGS, the *NC* approximation and its modification by CC considerably underpredict the effects of sweeps compared with simulations, whereas the *C2* approximation derived here fits much better. The *C1* approximation greatly overestimates the diversity reductions at high recombination rates.

This inaccuracy of the *NC* approximation reflects the fact that the probability of no recombination in the absence of coalescence (*P_nr_*) used in the *NC* approximation (Equation 11b) declines much faster with increasing recombination rate than does the true probability of no recombination, (1 – *P_r_*) (see Table 1 and Table S1 of File S1). The theory for large *r*/*s* is, however, not entirely satisfactory, as the *C2* approximation derived here uses a heuristic approach to modeling the effects of multiple recombination events, while the *C1* approximation ignores these events. Tables 1 and S1 show that the probability of a single recombination event is often less than half the net probability of a recombination event when *r*/*s* ≥ 0.08, so that multiple events cannot then be ignored. As described above, the true expected reduction in diversity probably lies between the *C1* and *C2* predictions. Ideally, both multiple recombination events and within-sweep coalescent events should be included in the model without the approximations used here. This was done by Kaplan *et al*. (1989), but no simple formula can be obtained by this approach.

Other stochastic treatments of the mean coalescent time associated with a sweep that should, in principle, allow for multiple recombination events and coalescence within the sweep, have been given by Stephan *et al*. (1992) and Barton (1998) for the case of semi-dominant selection with autosomal inheritance and random mating. However, these are not necessarily very accurate. For the example in Figure 1 with *h* = 0.5, *γ* = 250 and *r*/*s* = 0.64, Equation 18 of Stephan *et al*. (1992) predicts a reduction in diversity at the end of a sweep of 0.013, whereas the simulations and *C2* approximation give values of 0.0051 and 0.0044, respectively. After some simplification (and equating Barton’s *ε* to *q*_0_), Equation 16 of Barton (1998) gives a predicted reduction in pairwise diversity in the absence of recombination of approximately 1 – *T_d_* – 2*γ*^−1^ ln(γ). For *γ* = 250 (with *T_d_* = 0.088), this is equal to 0.868, compared with the simulation and *C1*/*C2* values of 0.925, and the *NC* value of 1. In addition, Figure 3 of Barton (1998) shows a slightly faster than linear increase in mean coalescent time with increasing *r*/*s* for *r*/*s* < 0.5, in contrast to the approximately exponential decline in – Δ*π* seen in Figure S1, corresponding to a diminishing returns relation for *π*.

It is important to note that, even for recombination events within genes, relatively large *r*/*s* values are likely. For example, with the parameters used for Figure 3, *s* = 1.25 x 10^−4^ in a population with *N_e_* = 10^6^. With the standard sex-averaged autosomal CO rate for *D. melanogaster* of 1 x 10^−8^ per bp, but without gene conversion, the recombination rate between two sites 1kb apart is 1 x 10^−5^, so that *r*/*s* = 0.08. With gene conversion at an effective rate of initiation of 1 x 10^−8^ per bp and a mean tract length of 440bp, the recombination rate is 1.88 x 10^−5^, and *r*/*s* = 0.15. Figure 1 shows that, for *r*/*s* = 0.16, *h* = 0.5 and *γ* = 250, the predicted reduction in diversity at the end of sweep for *C2* is approximately 86% of that for *C1* and 114% of the simulation value (this is not significantly different from the *C2* result at the 1% level); the *NC* approximation predicts a reduction that is approximately 27% of the *C2* value.

Knowledge of the expected effects of multiple recombination events for large *r*/*s* is even more important for modeling recurrent sweep effects on intergenic sequences, which is needed for interpreting the observed pattern of increased intergenic sequence variability as a function of the distance from a gene in both mammals (Halligan *et al*. 2010; Hammer *et al*. 2010; Booker 2018); and *Drosophila* (Johri *et al*. 2020). An improved analytical treatment of this problem is desirable. At present, the use of the *C2* approximation seems to provide the best option for dealing with recurrent sweeps, other than by numerical solutions using the results of Kaplan *et al*. (1989) or simulations of the type performed by Messer and Petrov (2013) and Johri *et al*. (2020).

### Relations between synonymous site diversity and recombination rate

The main purpose of this paper is to explore some general principles rather than to attempt to fit models to data, but it is obviously of interest to examine the relations between the theoretical predictions for the *Drosophila* model of recurrent selective sweeps described above and the relevant empirical evidence. As Figure 3 shows, despite the caveats discussed about, the analytical results derived here for selective sweep effects using the *C2* approximation should provide better predictions concerning the relation between synonymous site diversity and local recombination rate than those discussed in CC. The basic expectation of a diminishing returns relation between synonymous site diversity and CO rate described in CC remains, however, unchanged.

Based on the empirical plots of this relationship provided in Campos *et al*. (2014), it was concluded by CC that the observed relation between synonymous *π* and CO rate in a Rwandan population of *D. melanogaster* was too steep and close to linear to be explained by models that include both selective sweep and BGS effects, or by either of these processes on their own. One possibility is that CO events are mutagenic, as indicated by recent studies of human *de novo* mutations (Halldorsson *et al*. 2019). This is, however, hard to reconcile with the lack of evidence for a correlation between silent site divergence and CO rate in *D. melanogaster*, outside the non-crossover genomic regions where divergence tends to be *higher* than average, presumably reflecting the effect of reduced *N_e_* due to selection at linked sites (Haddrill *et al*. 2007; Campos *et al*. 2014). This observation is, however, not conclusive, since the recombination landscape in *D. melanogaster* is substantially different from that in its close relative *D. simulans*, with less suppression of crossing over near telomeres and centromere (True *et al*. 1996), so that current estimates of CO rates may not reflect the evolutionarily significant values.

Another possibility is that the nearly linear relationships between described by Campos et *al.* (2014) are artefacts of their use of classical marker-based maps (Fiston-Lavier *et al*. 2010) or the Loess smoothing procedure applied to the 100kb window estimates of CO rates obtained by the SNP-based map of Comeron *et al*. (2012). Smoothing may cause relative low values of *π* associated with very high CO rates to be wrongly assigned to much lower CO rate, as noted by Castellano *et al*. (2016) in connection with estimates of the relation between the rate of adaptive evolution of protein sequences and the CO rate. A diminishing returns relation between non-coding site diversity for the Raleigh population of *D. melanogaster* and CO rate was found by Comeron (2014) when using the raw CO rate estimates; a similar pattern is seen in the Rwandan population (J.M. Comeron, personal communication). However, the use of the raw estimates is open to the objection that the extreme CO values may simply be artefactual, leading to a flatter relation between *π* and CO rate than truly exists. In addition, Comeron’s non-coding *π* values for the Rwandan population are substantially lower than the synonymous site values of Campos et *al.* (2014), similar to what has been found in studies of other populations (Andolfatto 2005; Haddrill *et al*. 2005), suggesting that the non-coding sites involve at least some sequences that are subject to selection. This could easily lead to a less than linear relation between *π* and CO rate. The question of the true empirical relationship between neutral or nearly neutral variability in *Drosophila* needs further exploration before firm conclusions can be drawn.

### Differences between X chromosomes and autosomes

As discussed by CC, the differences between X chromosomes and autosomes in their levels of neutral diversity, and the relations between these and CO rates, need to be interpreted in terms of models of the effects of selection at linked sites. A pattern seen in several analyses of *D. melanogaster* datasets is that the relation between silent or synonymous site diversity for the X and CO rate is considerably weaker than that for the autosomes (Langley *et al*. 2012; Campos *et al*. 2014; Comeron 2014).

The expected difference between X and A is seen mostly clearly by plotting the ratio of X to A diversity values (*R_XA_*) against the CO rate, adjusted to give the same effective rate for X and A genes. The results for the case when *R_XA_* in the absence of selection (*k*) is equal to 0.75 were shown in Figure 6). The contrast between the cases with and without BGS is striking. Without BGS, for each mode of selection the ratio either slightly increases with CO rate (*h* = 0.1) or is constant or nearly constant (*h* = 0.5 or *h* = 0.9). With BGS, there is a strong decline in *R_XA_* from the lowest relative rate of CO (0.5), with values greater than 1 or close to 1, and the standard rate (1.0). The value at the highest relative CO rate (2.5) varies according to the dominance coefficient and mode of selection. With male-only selection, *R_XA_* ≈ 0.7 with *h* = 0.1 but is close to 0.8 with the other dominance coefficients; for the other modes of selection it is close to 0.75.

Given the evidence that the X in *Drosophila* is deficient in genes with male-biased expression, but enriched in female-biased genes (Parsch and Ellegren 2013), the results for male-biased genes are probably the least relevant. The data on synonymous site diversity in the Rwandan population of *D. melanogaster* in Figure S2 of Campos *et al*. (2014), based on Loess smoothing of the raw recombination estimates of Comeron *et al*. (2012), show that *R_XA_* takes the values of 1, 0.84. 0.74 and 0.73 for relative CO rates of 0.5, 1.0, 1.5 and 2.0, respectively. Use of the raw estimates of CO rates and diversity at non-coding sites for the same population gives a qualitatively similar pattern (J.M. Comeron, personal communication).

### Effect of a change in population size on *R_XA_*

It has been shown previously that a change in population size can cause *R_XA_* to deviate from its equilbrium value, *k* (Hutter *et al*. 2007; Pool and Nielsen 2007, 2008), reflecting the fact that the rate of response of neutral diversity to a change in population size is faster with smaller *N_e_*. This raises the question as to whether the observed pattern of relationship between *R_XA_* and the CO rate that has just been discussed could be explained by such a change, rather than by the differential effect of selection at linked sites on X and A diversity values. An approximate answer to this question can be obtained with a purely neutral model, in which the population size changes from a initial equilibrium value, but *k* remains constant during the process of change. In addition to *k*, we need to specify the relation between the rate of recombination and diversity. This can be done by introducing a variable *β* (0 ≤ *β* ≤ 1) which is equal to the ratio of the equilibrium diversity for a given effective rate of recombination to its value at the maximum recombination rate in the study. On the null hypothesis that there is no differential effect on *R_XA_* of selection at linked sites, the same *β* should apply to X and A diversities with the same effective recombination rate.

The most extreme effect of a population size change on *R_XA_* will come from a step change in population size, since this minimizes the ability of diversity values to track the population size. Consider a model in which the time *T* since the start of the expansion is scaled relative to the final autosomal *N_e_* at the highest recombination rate; let the ratio of final to initial effective population sizes be *R_N_*; and write *α* = 1 – 1/*R_N_*. Using the equivalents of Equation A8a applied to X and A diversities relative to their final equilibrium values, *R_XA_* at time *T* is given by:

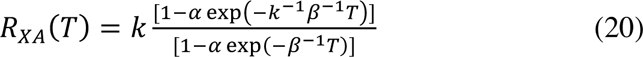

This expression shows that, as expected, *R_XA_* is equal to *k* when *T* = 0, and also when *T* >> *kβ*. For a population expansion (so that 0 < *α* < 1), for intermediate *T* values we have *R_XA_* > *k*, provided that *k* < 1. The converse is true for a population contraction, for which *α* < 0. Thus, regardless of the value of *β*, a population expansion will temporarily increase *R_XA_* above its equilibrium value. The extent of this increase for a given *T* is affected by *β*, but the direction and magnitude of the effect of *β* varies with *T*. For small *T*, a smaller value of *β* causes a larger value of *R_XA_*; the reverse is true for large *T* (see Section 1 of File S1).

It follows that no simple predictions are possible as to whether low rates of recombination are associated with larger values of *R_XA_* than high rates, after a population expansion of the kind indicated by the data on the Rwandan population of *D. melanogaster*. However, numerical examples suggest that the effects of differences in *β* are at best modest when *k* = 0.75. For example, with *R_N_* = 10 (a ten-fold increase in population size), the values of *R_XA_* for *β* = 0.25 and *β* = 1 are 0.892 and 0.858, respectively at *T* = 0.1. By *T* = 0.2, the respective *R_XA_* values are 0.869 and 0.885, and by *T* = 0.5 they are 0.801 and 0.884. There is therefore only a brief interval of time in which the lower *β* value (which corresponds to the lowest recombination rate considered above) is associated with *R_XA_* substantially larger than that with the higher value (which corresponds to the higher recombination rate considered). With *β* = 1, *R_XA_* > 0.85 persists until*T* = 1.

Instead of comparing X and A, we can compare the ratio of diversity values for two different regions of the same chromosome with different CO rates, with the left-hand side of Equation 20 representing this ratio at a given time after a population size change, where *k* now represents the effect of recombination rate differences on the equilibrium level of diversity (*β* is set equal to 1, since we are now longer comparing X and A). Equation 20 shows that a population expansion will reduce the differentials between regions with different *k* values, whereas a contraction will enhance them. For example, with *R_N_* = 10, *k* = 0.5 and *T* = 0.1, the diversity ratio becomes 0.709 instead of 0.5. Given the distortion of the site frequency spectrum at synonymous sites on the autosomes in the Rwandan population towards low frequency variants, with Tajima’s *D* values at synonymous sites of approximately – 0.2 (Campos *et al*. 2014), there has probably been a recent population expansion, which may have weakened the relation between diversity and CO rate compared with an equilibrium population. Further theoretical investigations of the interaction between such demographic effects and effects of selection at linked sites are needed if reliable inferences concerning both demography and selection are to be obtained (Messer and Petrov 2013; Zeng 2013; Ewing and Jensen 2016; Comeron 2017; Becher *et al*. 2020; Johri *et al*. 2020).

## Acknowledgments

I thank Matthew Hartfield for many useful discussions of this problem and for supplying me with the simulation results used in Figures 2 and S2. I also thank Josep Comeron for helpful discussions, and for sharing his analyses of Drosophila polymorphism data. I thank Nick Barton, Matthew Hartfield and an anonymous reviewer for their comments on the original manuscript, which led to significant improvements.

## Appendix

### Explicit formulae for coalescence and recombination probabilities

When *a* > 0 and *a* + *b* > 0 (i.e. excluding cases of random mating with complete recessivity or dominance), the time between a given frequency *q* of A_2_ and its frequency *q*_2_ at the end of the deterministic phase of the sweep is given by:

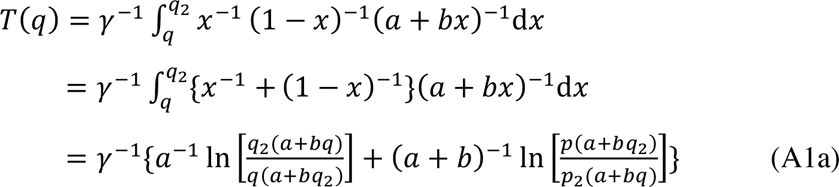

When *a* tends to 0 and *b* tends to 1, corresponding to random mating with complete recessivity, we have:

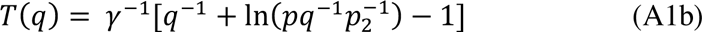

When *a* tends to 1 and *b* tends to –1, corresponding to random mating with complete dominance, we have:

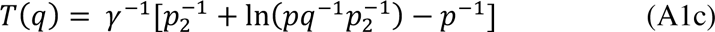

Similarly, for *a* > 0 and *a* + *b* > 0 Equation 8b can be written as:

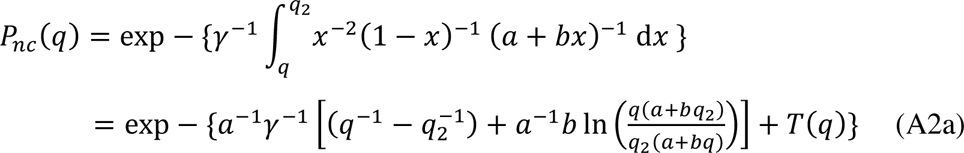

When *a* tends to 0 and *b* tends to 1 (complete recessivity), we have:

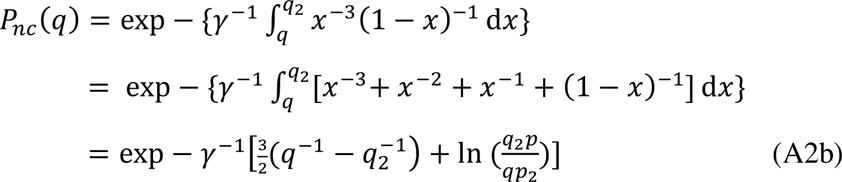

When *a* tends to 1 and *b* tends to –1 (complete dominance, we have:

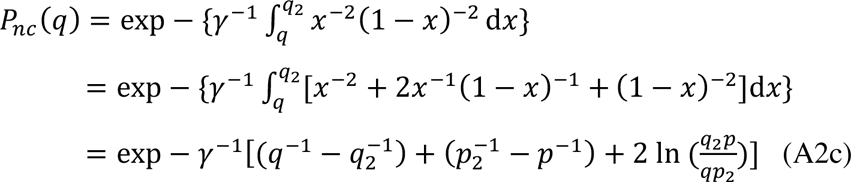

Equation 8c for *a* > 0 and *a* + *b* > 0 can be written as:

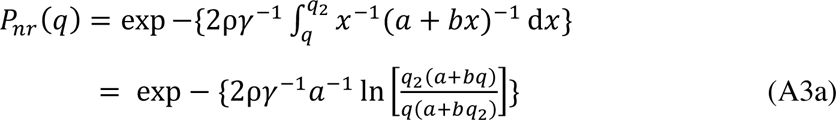

When *a* tends to 0 and *b* tends to 1, this becomes:

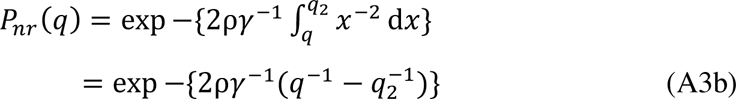

When *a* tends to 1 and *b* tends to –1, we have:

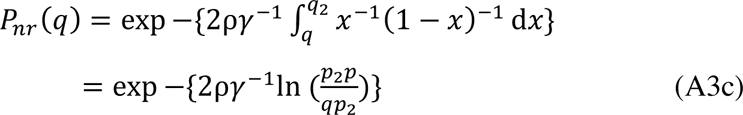

### Probability of no coalescence conditional on no recombination

For simplicity, only the case of intermediate dominance (*a* ≠ 0, *b* ≠ –1) will be considered. With large *γ*, we can write *q*_1_ ≈ 1/(2*aγ*), *p*_2_ ≈ 1/[2(*a* + *b*)*γ*] and *q*_2_ = *p*_1_ ≈ 1. We can then use Equations A1a and A2a with *q* = *q*_1_. *T*(*q*_1_) and the multiplicand of *b* in the exponent are of order ln(*γ*)/*γ*, provided that *a*^−2^ << *γ*. The leading term in the exponent is the product of – 1/(2*aγ*) and 1/*q*_1_, which is approximately equal to –2 under this condition, implying that *P_nc_*(*q*_1_) ≈ *e* ^−2^ = 0.135.

### Harmonic mean of *q* during a sweep

The integral of 1/*q* between *q*_1_ and *q*_2_ is equivalent to the terms in braces in the exponents in the first lines of Equations A3, with *q* = *q*_1_. The harmonic mean of *q* is given by taking the reciprocal of this integral, after division by *T_d_*. From the above result, the integral is approximately 2 for large *γ*, so that the harmonic mean of *q* is approximately equal to ½*T_d_*.

### Probability of recombination during a sweep, for large values of *r*/*s*

A semi-dominant (*h* = 0.5) autosomal locus with random mating is assumed. Using the notation described in the main text, together with Equations A1a, A2a, and A3a, Equation 10 can be written as:

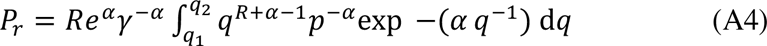

Assuming that *γ* >> 1 (so that *α* << 1), and *R* > 1, the integrand can be approximated by:

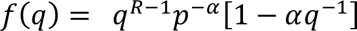

Expanding *p*^−^*^α^* as a binomial series in powers of *q*, the integrand can be written as:

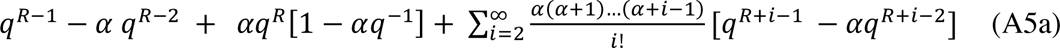

Neglecting higher powers in *α*, this can be approximated by:

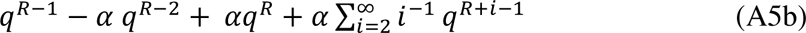

This expression can be integrated term by term to yield the following indefinite integral:

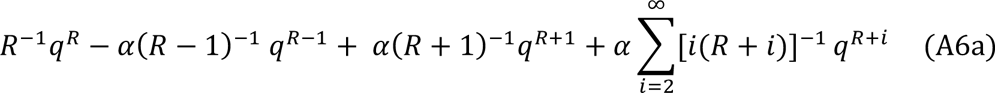

We can then use *q*_1_ = *p*_2_ = 1/*γ* to obtain the definite integral in Equation A4, again neglecting higher order terms in *α*:

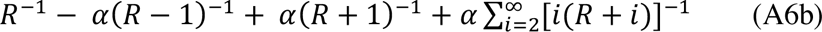

Simplifying the first two terms in *α* in this expression, and multiplying by the terms outside the integral in Equation A4, we obtain the following expression for the probability of recombination during a sweep:

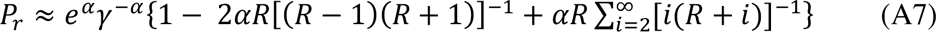

### Increase in expected diversity of recombinant alleles at the start of a sweep

At the start of a sweep, the expected diversity (relative to the neutral expectation) is *π*_1_. If there is sufficient recombination that recombinant alleles have an effective population size of *B*_1_ relative to neutrality, their expected relative pairwise diversity relative to neutrality at a given time *T* since the start of the sweep is given by:

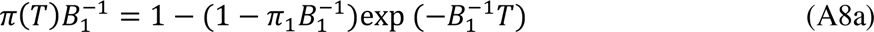

(Compare with Equation 11 of CC).

For sweep of type *i,* with duration *T_di_* and mean time to the first recombination event *T_ri_*, the expected diversity at the time of the recombination event is thus:

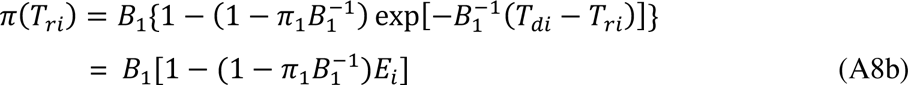

If we assume that this is approximately the same as the mean coalescent time associated with multiply-recombinant haplotypes, the diversity at the end of a sweep of this type can be written as:

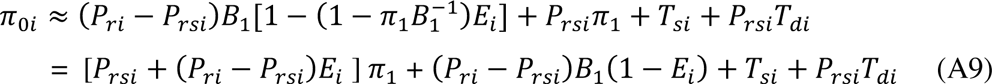

If the rate of occurrence of sweeps of type *i* is *ω_i_*, their frequency among all sweeps is equal to *f_i_* = *ω_i_* /*ω*. The expected diversity at the end of a sweep is given by:

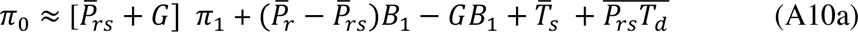

Where

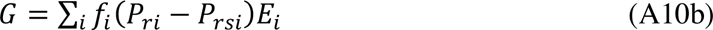

From Equation 10 of CC, we also have:

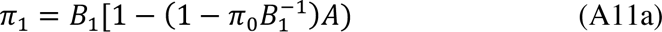

Where

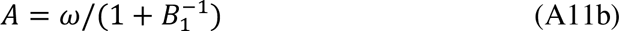

Furthermore, by Equation 10 of CC, the mean diversity over the interval between sweeps, *π*, is given by:

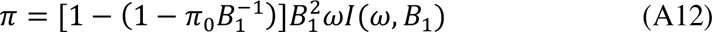

where *I* is the integral defined in Equation 10 and section 5 of File S1of CC.

As in CC, these expressions allow *π*_0_, *π*_1_ and *π* to be determined explicitly. Let 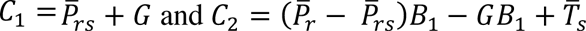. Using Equations A10 and A11, we have:

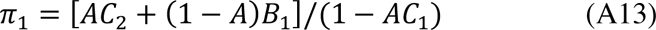

Substituting this expression into Equation A10a yields an expression for *π*_0_, which in turn allows *π* to be determined from Equation A12.

### Rates of substitution of new favorable mutations for arbitrary genetic systems

Following Charlesworth *et al*. (2018), let *N_H_* be the total number of haploid copies among breeding adults for a given genetic system, where *N_H_* = 2*N* for A, *N_H_* = 3*N*/2 for X, and *N* is the number of breeding adults. Let the effective population size for the genetic system in question be *kN_e_*_0_, where *N_e_*_0_ is the effective population size for A. Under the selection model described in the text (Equation 6), the probability of fixation of a new mutation is 4*as*(*kN*_e0_/*N_H_*) (Charlesworth 2020), and the rate of entry of new favorable mutations into the population each generation is *N_H_up_a_,* where *p_a_* is the proportion of mutations that are favourable. The rate of substitution of favorable mutations per generation is thus 4*N_H_up_a_* (*as kN_e_*_0_/*N_H_*) = 4*kN_e_*_0_*up_a_* (*as*) = 2*up_a_* (*akγ*_0_), where *γ*_0_ is the scaled selection coefficient for autosomal mutations. The expected number of substitutions over one unit of coalescent time for the given genetic system is thus 4*kN_e_*_0_ x *up_a_* (*akγ*_0_) = *πp_a_*(*akγ*_0_), where *π* = 4*kN_e_*_0_*u* is the expected neutral nucleotide site site diversity with mutation rate *u*. The relevant formulae for *a* described in the text can be used to obtain the substitution rates for the genetic system of interest.

## Section S1

### Derivation of Frisse et al. formula for the effect of gene conversion

Consider two sites (1 and 2, with 1 to the left of 2) separated by *z* basepairs; let the rate of initiation of a gene conversion tract be *r_g_* and the mean tract length be *d_g_*. Assume an exponential distribution of tract lengths, with rate parameter *λ* = 1/*d_g_*. If a tract is initiated to the left of 1, it has a probability of ½ of moving towards 1. If it is initiated at distance *y* from 1, the probability that it includes 1 but falls short of 2, resulting in the conversion of 1 but not 2, is given by:

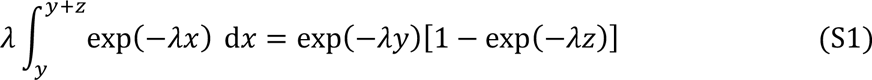

The net probability of conversion of 1 from this class of event is then given by:

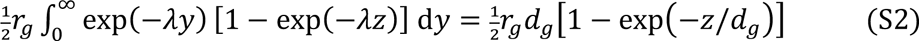

We also need to consider the class of events where a tract is initiated between sites 1 and moves the right including 2. If it starts at a distance *u* from 1, the probability that it includes 2 is given by:

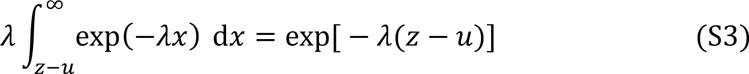

The net probability of a tract of this class converting site 2 but not 1 is:

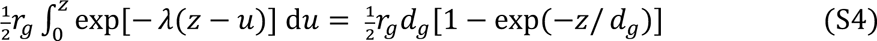

The same argument applies to tracts moving from right to left, so the net probability of recombination between sites 1 and 2, due to one site but not the other being included in a conversion tract, is:

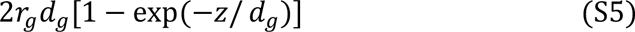

### Effect of the *β* parameter on *R_XA_*

It is sufficient to consider the partial derivative of *R_XA_* with respect to 1/*β*, as given by Equation 19 of the main text. We have:

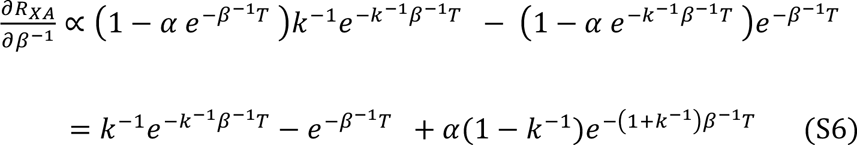

As *T* tends to 0, this expression approaches (1 – *α*)(*k* ^−1^ – 1). Provided that *k* < 1, this is positive if *α* < 1 (a population expansion), and negative if *α* > 1 (a population contraction). The product of exp(*βT*) and the derivative is equal to:

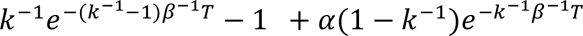

For *T* >> *kβ* and *k* < 1, this quantity approaches –1, which implies that the derivative must be negative; a negative value will be reached at smaller values of *T*, the larger *α*. In the case of a population expansion, this corresponds to a smaller increase in population size. The effect of *β* on *R_XA_* for a given value of *k* is therefore dependent on the time since the start of the expansion.

## Section S2

**Table S1.**
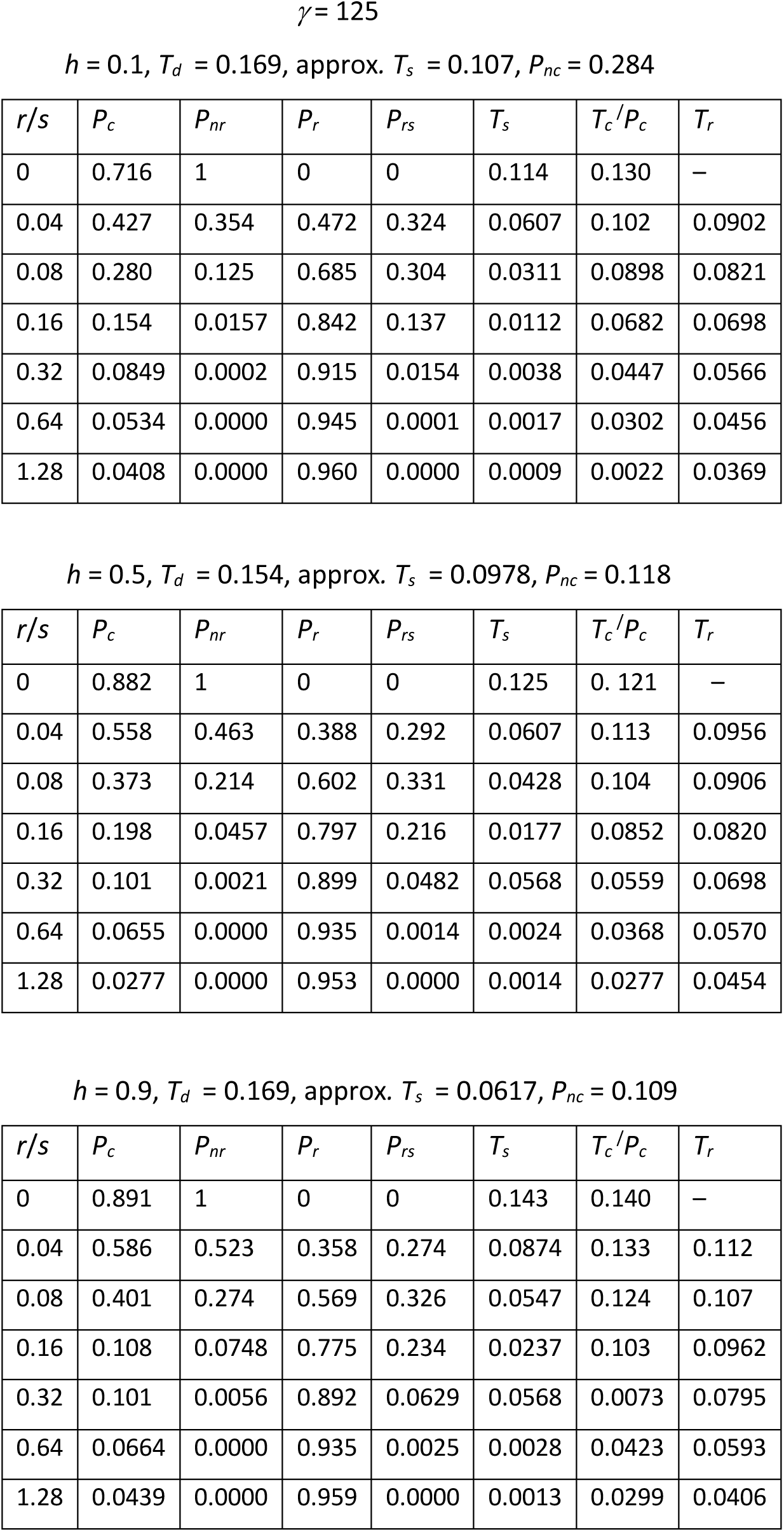
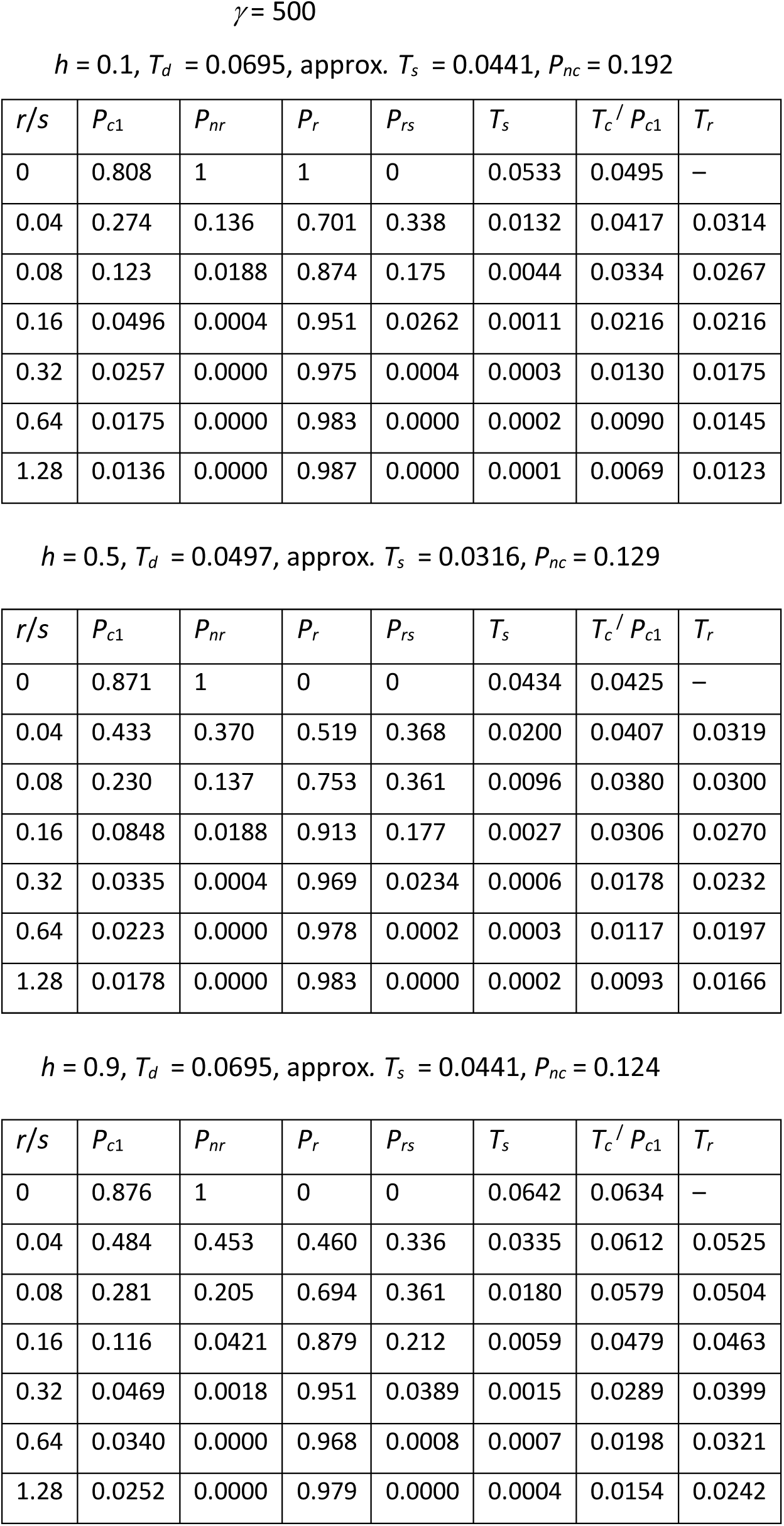
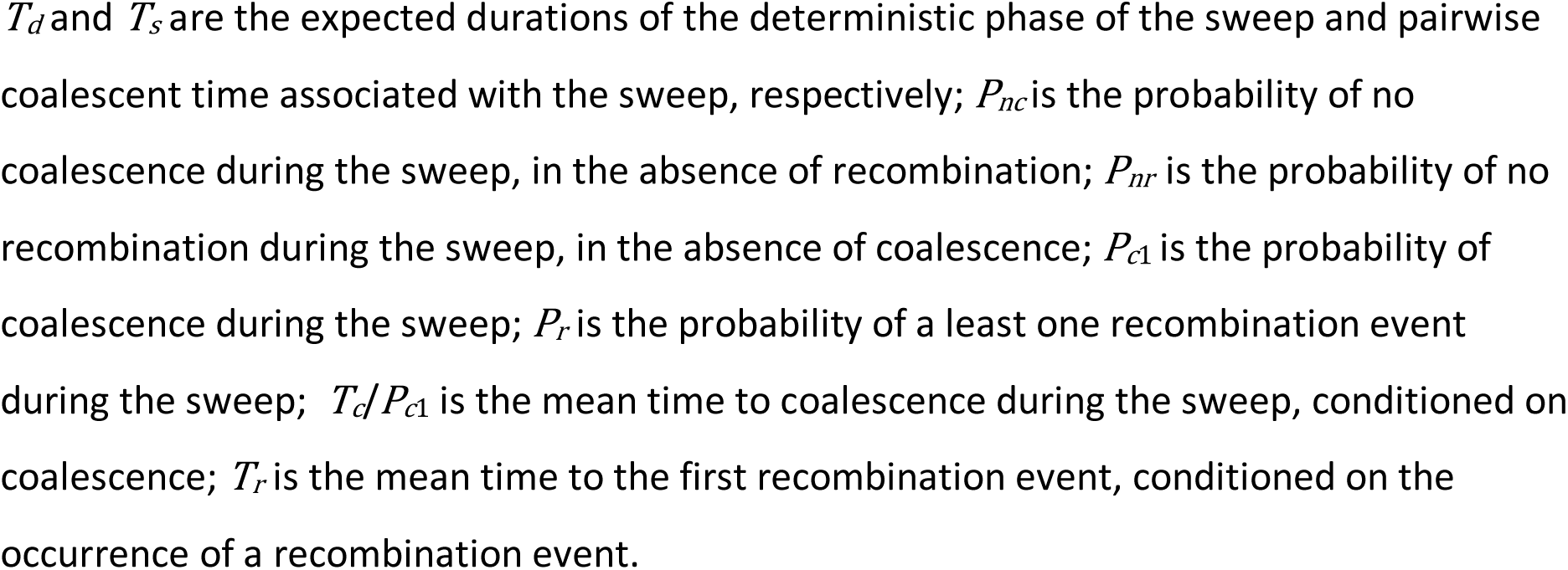
**Results for a single sweep of an autosomal locus in a randomly mating population with N = 106**

## Section S3

**Figure S1.**
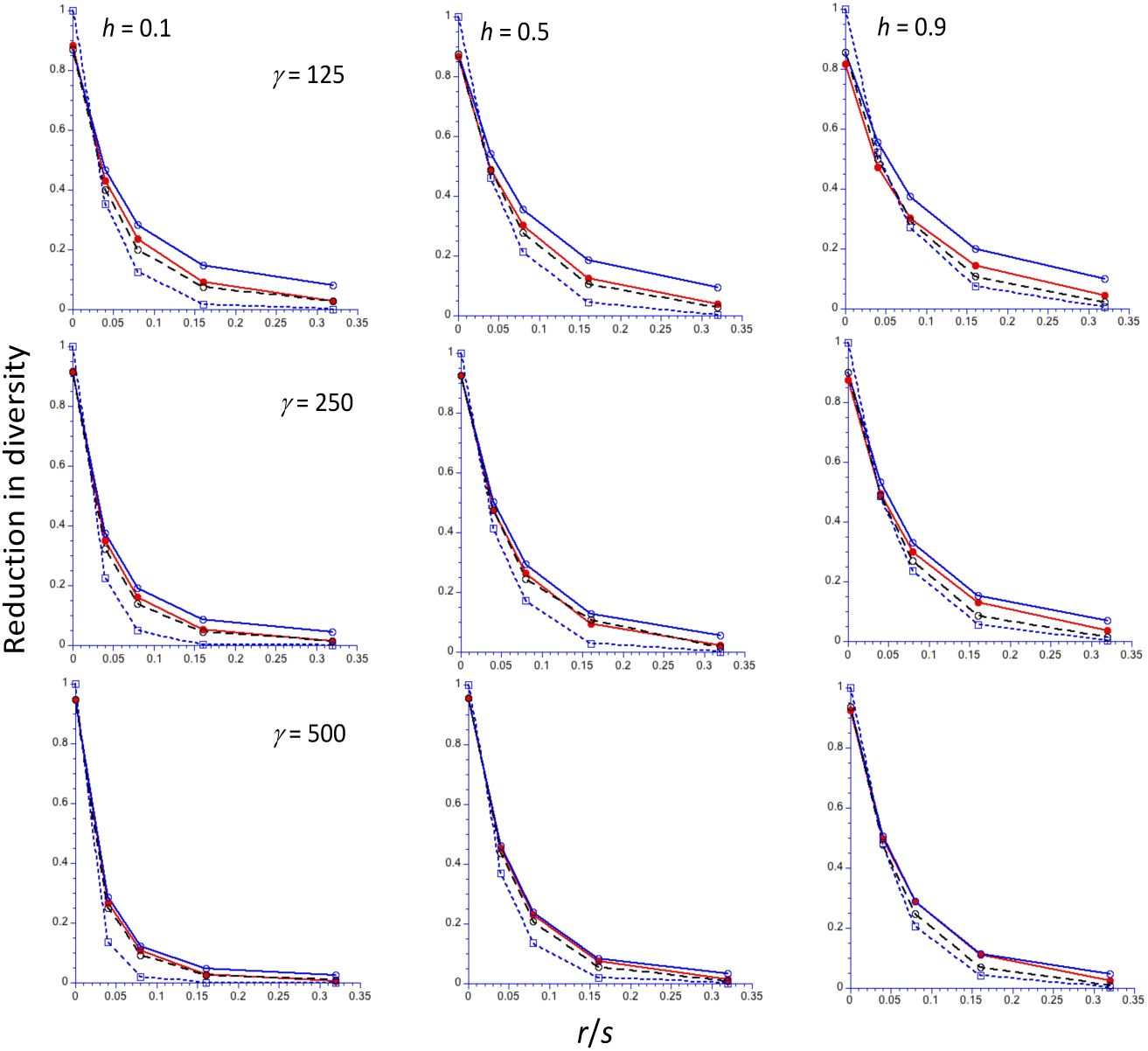
The reduction in diversity (relative to the neutral value) at the end of a sweep for an autosomal locus (Y-axis, linear), as a function of the ratio of the frequency of recombination (*r*) to the selection coefficient for homozygotes (*s*) (X-axis, linear scale). A randomly mating population of size 5000 is assumed, with three different values of the scaled selection coefficient (*γ* = 2*N_e_s*): 125 (top panel), 250 (middle panel) and 500 (bottom panel), and three different values of the dominance coefficient (*h*), increasing from left to right. The filled red circles are the mean values from computer simulations, using the algorithm of Tajima (1990); the open blue circles and black circles are the *C1* and *C2* predictions, respectively; the open blue squares are the *NC* predictions.

**Figure S2.**
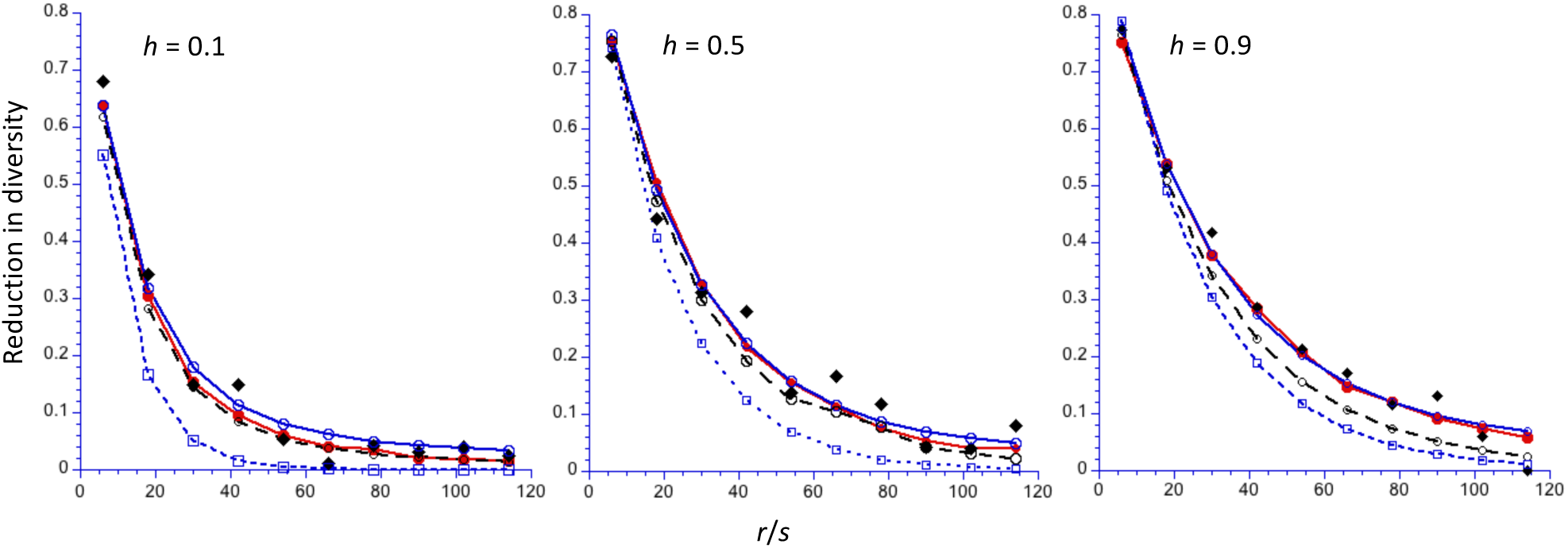
The reduction in diversity (relative to the neutral value) at the end of a sweep for an autosomal locus, as a function of the scaled rate of recombination (2*N_e_r*). The results for three different values of the dominance coefficient (*h*) are displayed. A randomly mating population of size 5000 is assumed, with a scaled selection coefficient (*γ* = 2*N_e_s*) of 500. The filled red circles and black lozenges are the mean values from computer simulations, using the algorithm of Tajima (1990) and the results of Hartfield and Bataillon (2020), respectively; the open blue circles and black circles are he *C1* and *C2* predictions, respectively; the open blue squares are the *NC* predictions.

**Figure S3.**
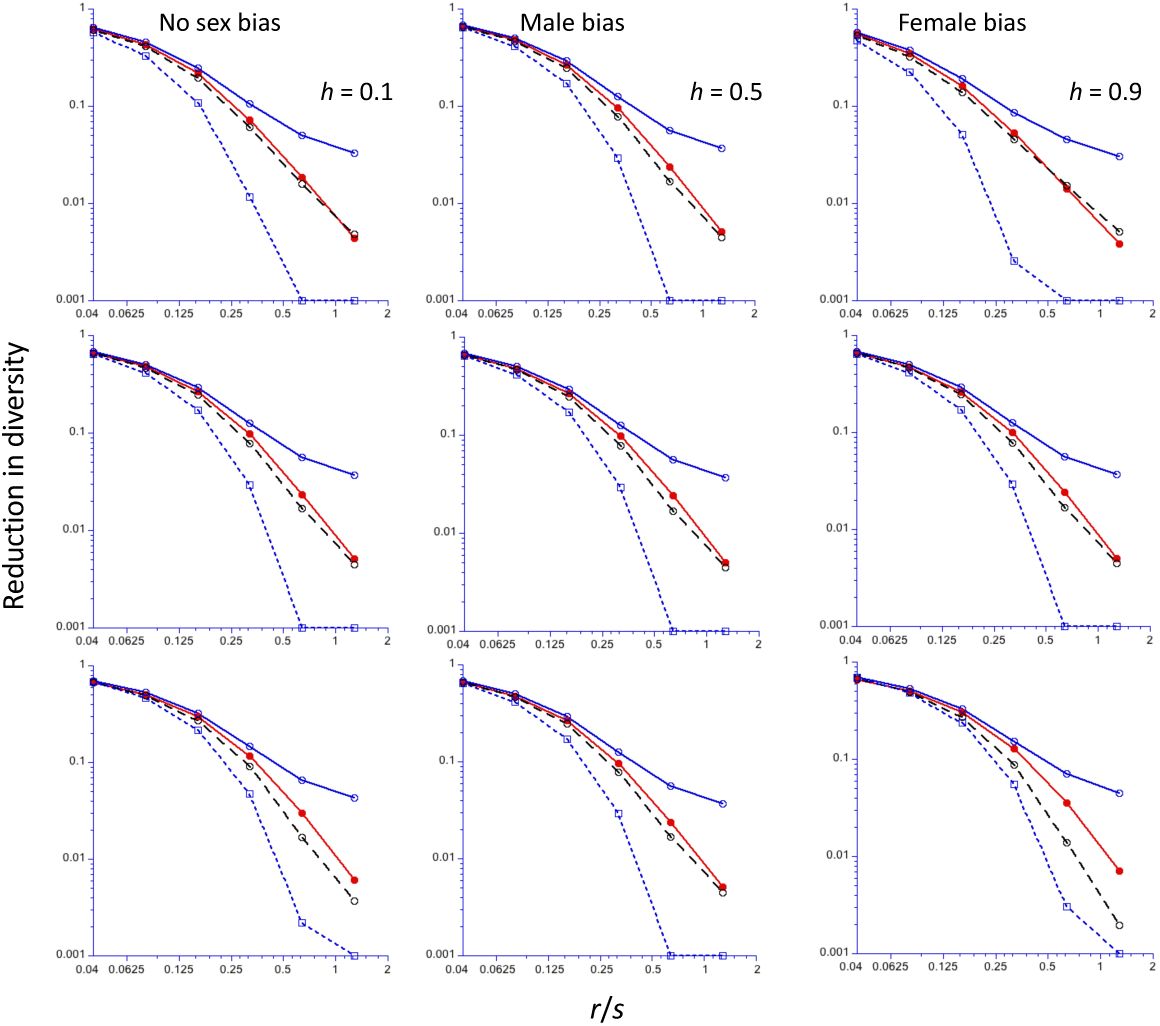
The reduction in diversity (relative to the neutral value) at the end of a sweep for an X-linked locus (Y-axis, log_10_ scale), as a function of the ratio of the frequency of recombination (*r*) to the selection coefficient for homozygotes (*s*) (X-axis, log_2_ scale). The mammalian recombination model is assumed; the The results for mutations with no sex limitation are shown in the left-hand panels; those for male-limited and female-limited mutations are shown in the middle and right-hand panels, respectively. A population size of 5000 is assumed, with a scaled selection coefficient for an autosomal mutation in a randomly mating population (*γ* = 2*N_e_s*) of 250 for the cases with no sex-limitation. For the sex-limited cases, *γ* = 500 to ensure comparability to sex-limited autosomal mutations. Results for three different values of the dominance coefficient (*h*) are shown, with *h* increasing from top to bottom. The filled red circles are the mean values from computer simulations, using the algorithm of Tajima (1990); the open filled blue circles and black circles are the *C1* and *C2* predictions, respectively; the open blue squares are the *NC* predictions. Values of the reduction in diversity less than 0.001 have been reset to 0.001.

**Figure S4.**
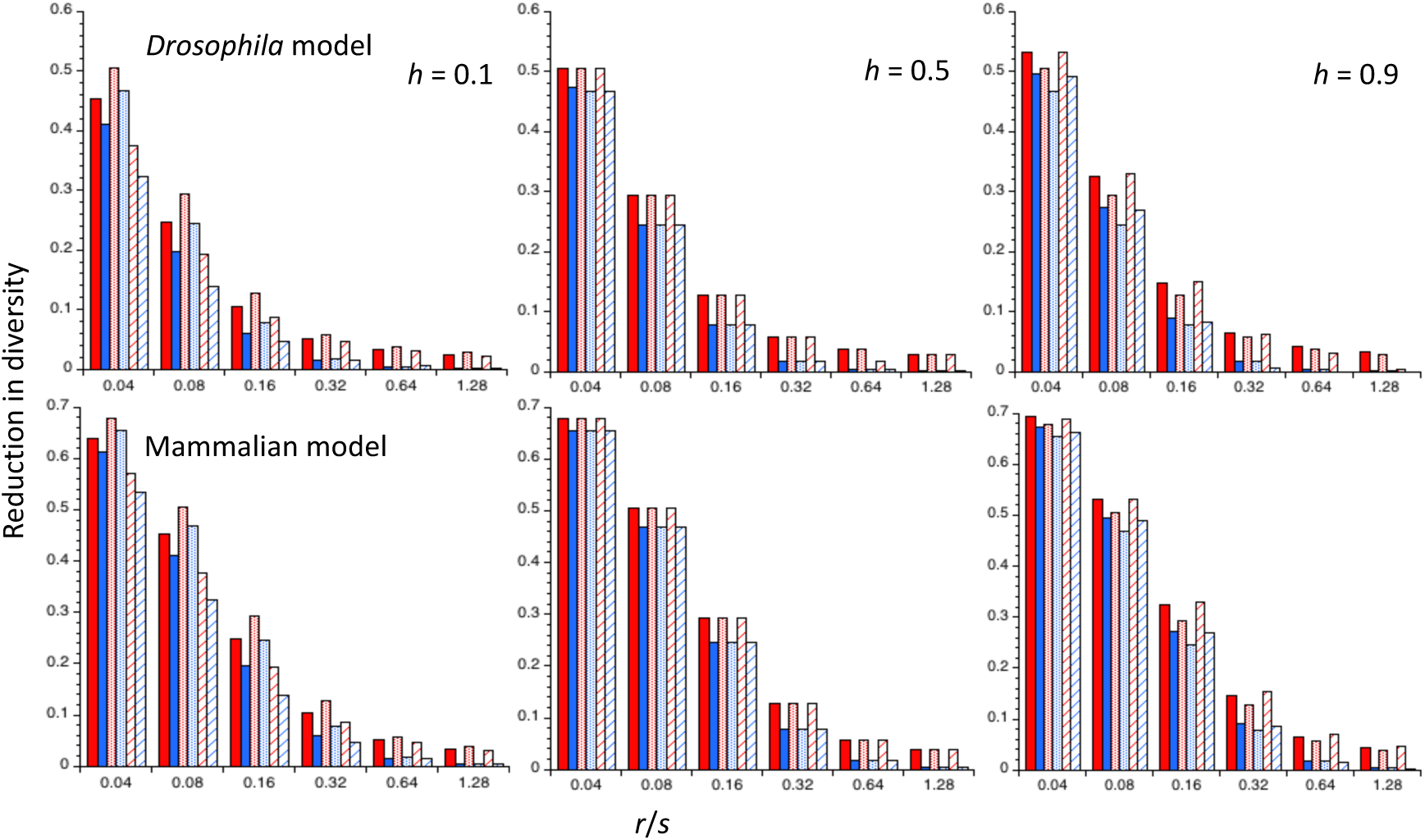
Reductions in diversity (relative to the neutral value) at the end of a sweep for an X-linked locus in a randomly mating population, as a function of the ratio of the autosomal value of the rate of crossing over to the homozygous selection coefficient (*r*/*s*), for three different dominance coefficients (*h*). *N_e_* for the X chromosome is three-quarters of that for the autosomes. Linear scales are used for both X and Y axes. The upper panel is for a *Drosophila* model, with no crossing over in males; the lower panel is for a mammalian model, with equal rates of crossing over for autosomes in both sexes. The red and blue colors denote the *C1* and *C2* predictions, respectively. The full bars denote results for mutations with equal effects in both sexes, with *γ* = 250. The stippled bars denote male-limited mutations and the hatched bars denote female limited mutations. For the sex-limited cases, *γ* = 500 to ensure comparability with sex-limited autosomal mutations.

**Figure S5.**
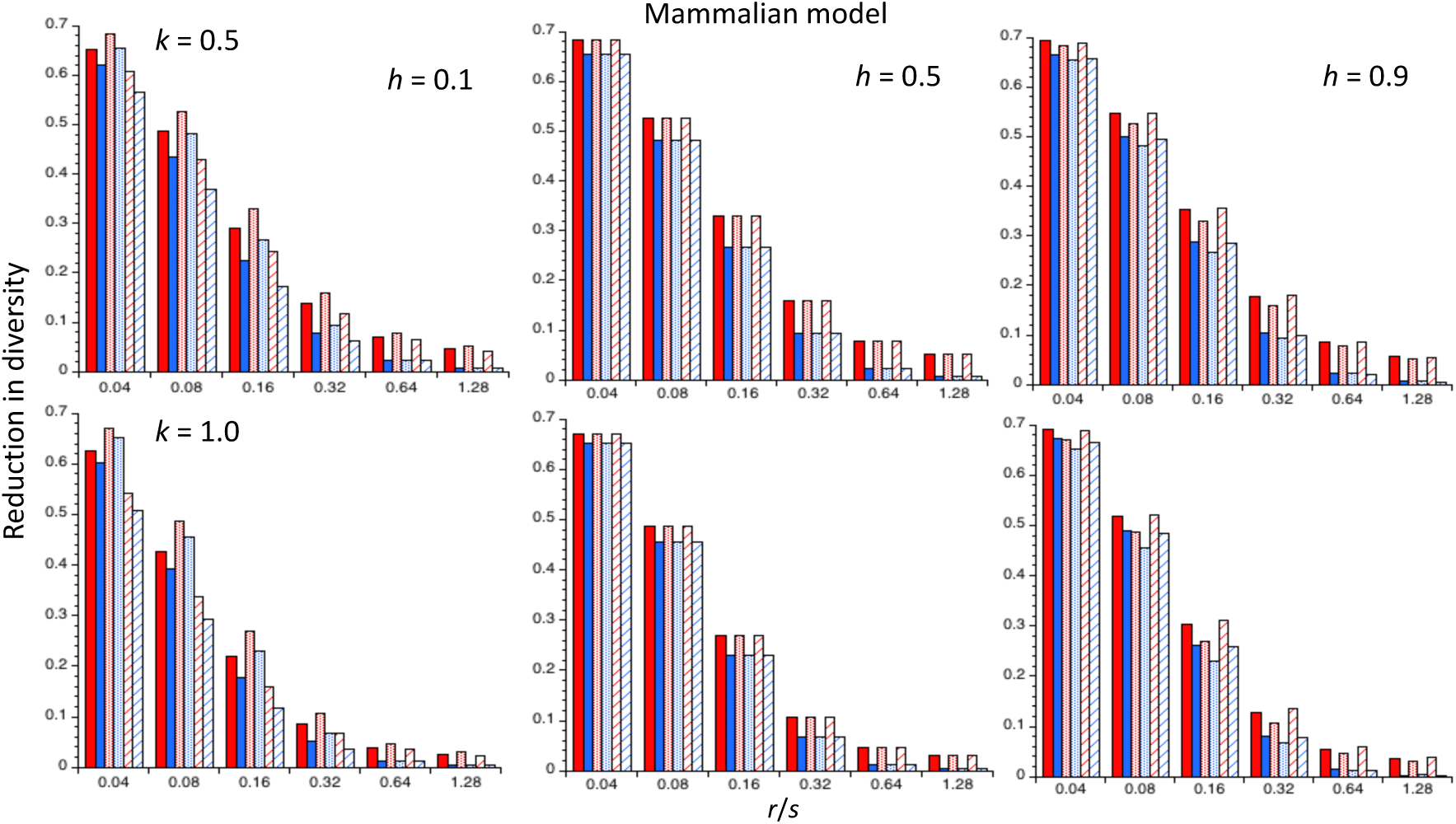
Reductions in diversity (relative to the neutral value) at the end of a sweep for an X-linked locus in a randomly mating population for the mammalian model (crossing over on the autosomes in both sexes), as a function of the ratio of the autosomal value of the rate of crossing over to the homozygous selection coefficient (*r*/*s*), for three different dominance coefficients (*h*). Gene conversion is absent. The upper panel is for the case when *N_e_* for the X chromosome is half of that for the autosomes; in the lower panel, *N_e_* is the same for both X and A. The red and blue colors denote the *C1* and *C2* predictions, respectively. The full bars denote results for mutations with equal effects in both sexes, with *γ* = 250. The stippled bars denote male-limited mutations, and the hatched bars denote female-limited mutations. For the sex-limited cases, *γ* = 500 to ensure comparability with sex-limited autosomal mutations.

**Figure S6.**
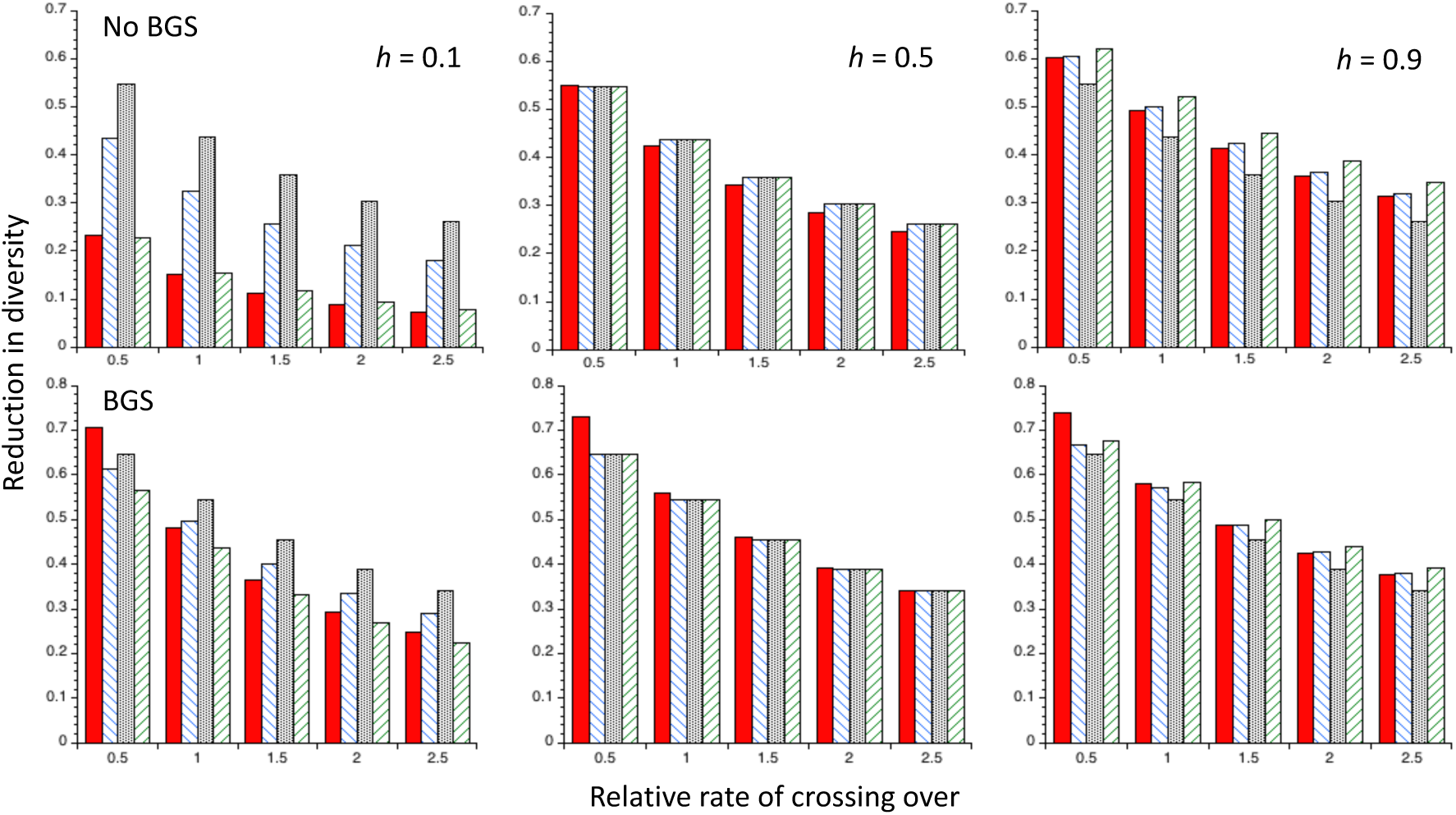
Reductions in diversity (relative to neutrality) under recurrent sweeps at autosomal and X-linked loci for the *Drosophila* model, using the *C2* theoretical predictions with no gene conversion and five different rates of crossing over relative to the autosomal standard value (the X-linked rates of crossing over were chosen to give the same sex-averaged effective recombination rates as for autosomes). *N_e_* for the X chromosome is three-quarters of that for the autosomes. The upper panel is for cases without BGS; the lower panel is for cases with BGS (using the parameters described in the main text). The filled red bars are for autosomal mutations, the hatched blue bars are for X-linked mutations with no sex-limitation, the stippled black bars are for male-limited mutations X-linked mutations, and the hatched green bars are for female-limited X-linked mutations. For the sex-limited cases, *γ* = 500 to ensure comparability with sex-limited autosomal mutations.

**Figure S7.**
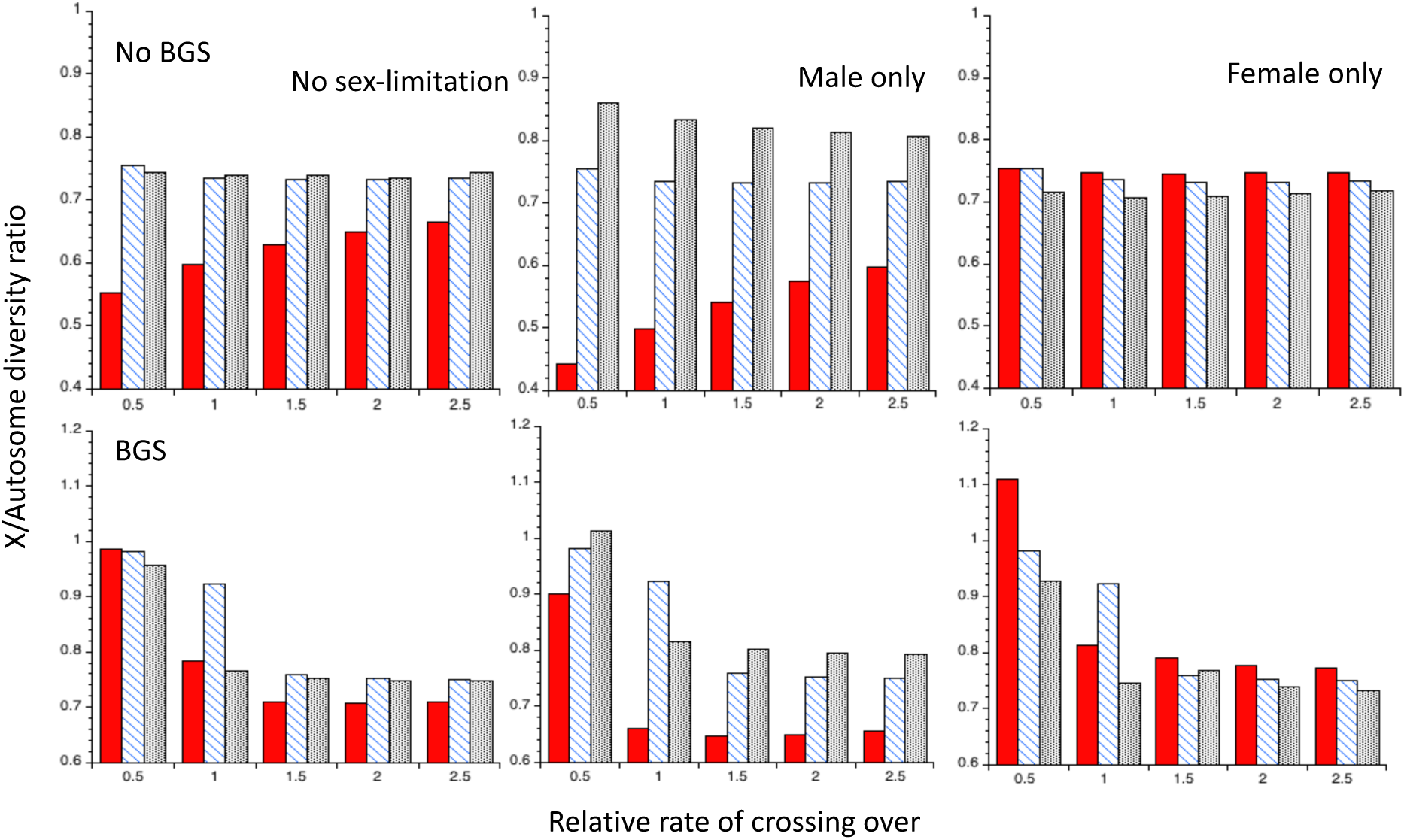
The ratios of X chromosome to autosome nucleotide site diversities under recurrent sweeps for the *Drosophila* model, using the *C2* theoretical predictions with no gene conversion and five different rates of crossing over relative to the autosomal standard value (the X-linked rates of crossing over were chosen to give the same sex-averaged effective rates as for the autosomes). The upper panel is for cases without BGS; the lower panel is for cases with BGS. The filled red bars are for *h* = 0.1 (the dominance coefficient of favorable mutations), the hatched blue bars are for *h* = 0.5 and the black stippled bars are for *h* = 0.9. The other details are as for Figure S6.

**Figure S8.**
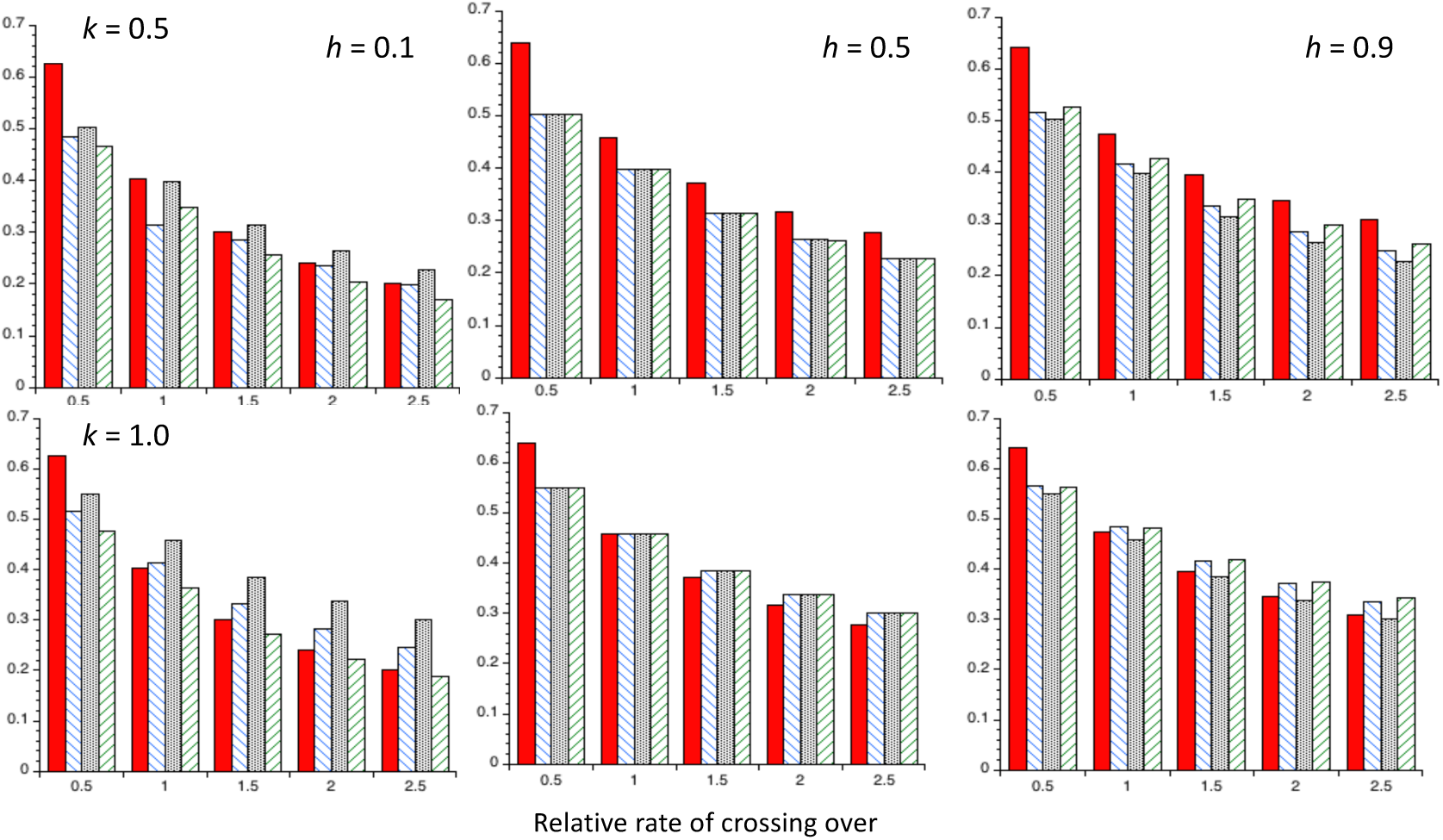
Reductions in diversity (relative to neutrality) under recurrent sweeps at autosomal and X-linked loci for the *Drosophila* model, using the *C2* theoretical predictions with gene conversion and five different rates of crossing over relative to the autosomal standard value (the X-linked rates of crossing over and gene conversion were chosen to give the same sex-averaged effective rates as for the autosomes). BGS is assumed to be present, with the parameters described in the main text. The upper panel is for the case when *N_e_* for the X chromosome is half of that for the autosomes; in the lower panel, *N_e_* is the same for both X and A. The filled red bars are for autosomal mutations, the hatched blue bars are for X-linked mutations with no sex-limitation, the stippled black bars are for male-limited mutations, and the hatched green bars are for female-limited mutations. For the sex-limited cases, *γ* = 500 to ensure comparability with the autosomal and non-sex-limited X-linked mutations.

**Figure S9.**
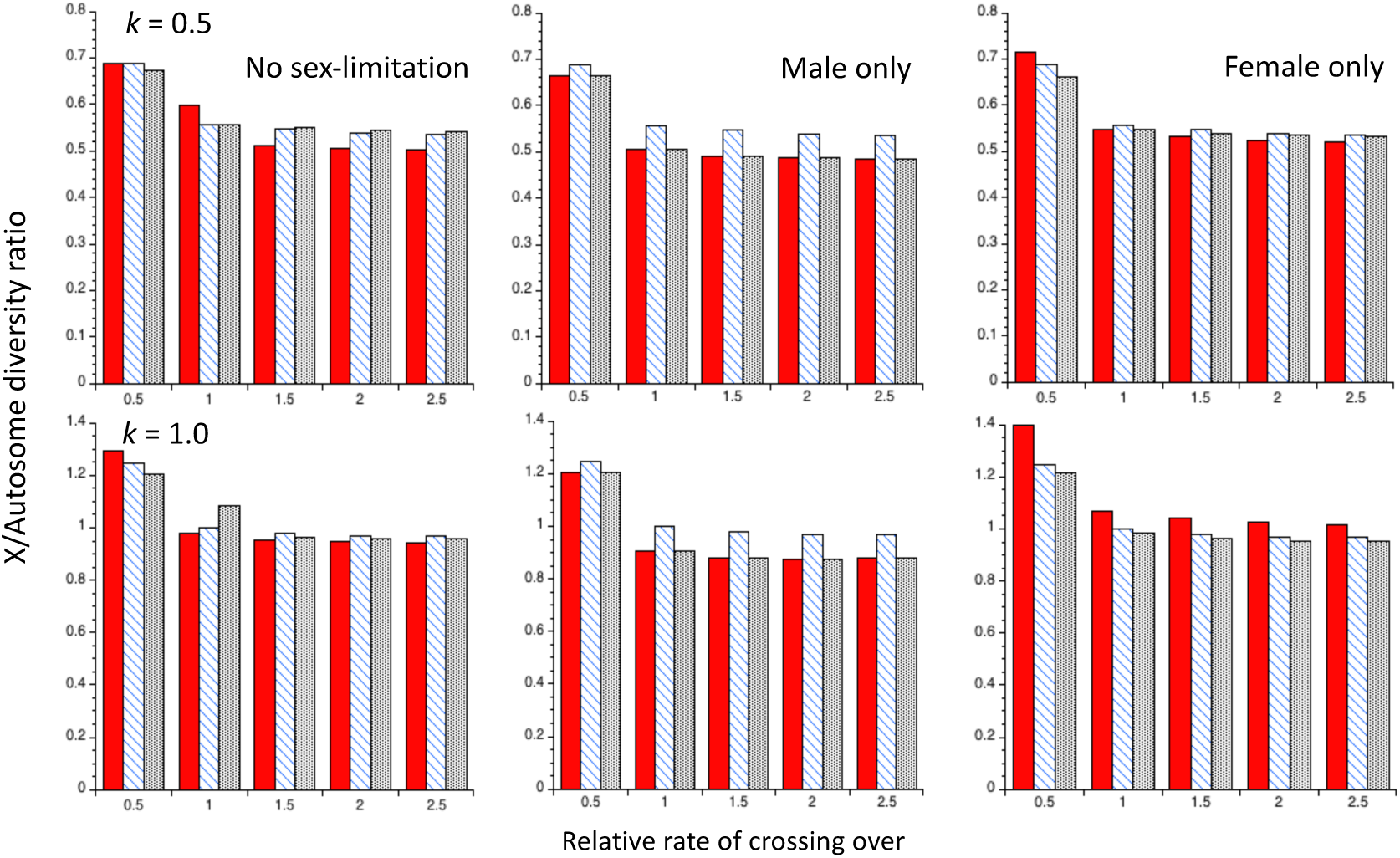
The ratios of X chromosome to autosome nucleotide site diversities for the *Drosophila* model with recurrent sweeps, using the *C2* theoretical predictions with gene conversion and five different rates of crossing over relative to the autosomal standard value (the X-linked rates of crossing over and gene conversion were chosen to give the same sex-averaged effective rates as the autosomal rates). BGS is assumed to be present. The upper panel is for the case when *N_e_* for the X chromosome is half that for the autosomes; in the lower panel, *N_e_* is the same for both X and A. The filled red bars are for *h* = 0.1 (the dominance coefficient of favorable mutations), the hatched blue bars are for *h* = 0.5 and the black stippled bars are for *h* = 0.9. The other details are as for Figure S8.

